# Silencing Proline Dehydrogenases Improves Salt and Drought Tolerance in Gossypium hirsutum

**DOI:** 10.1101/2025.07.02.662779

**Authors:** Suli Bai, Pei Zhao, Fei Su, Junjuan Wang, Shuai Wang, Yan Li, Ibrahim A. A. Mohamed, Zujun Yin

## Abstract

Proline is a key compound that lowers cell water potential, scavenges reactive oxygen species, and stabilizes biomolecules and cell membranes, thus reducing stress-induced damage. Proline dehydrogenase (ProDH), the first rate-limiting enzyme in proline degradation, plays a crucial role in proline accumulation. We explored the role of the GhProDH gene family in regulating physiological responses to stress conditions through transcriptome, metabolome and functional analyses. Overexpression and gene silencing lines in *Arabidopsis* and cotton revealed that *GhProDH2* plays a key role in regulating cotton’s tolerance to drought and salt stress. *GhProDH2-4* improves cotton’s tolerance to drought and salt stress by engaging in carbon metabolism, glyoxylate cycle, and flavonoid metabolism pathways. These findings highlight the potential role of *GhProDH2-4* to improve cotton stress resistance by genetically modifying pathways involved in proline biosynthesis and degradation.

## 1. Introduction

Drought and salt stress can cause water loss in plant cells, disrupting normal physiological processes. Salt stress disturbed ion toxicity (Hanin, Ebel et al. 2016) and nutrient imbalance in plant (Verma, Jalmi et al. 2020). Dehydration not only impairs plant growth and development but also directly reduces crop yield (Krasensky et al. 2012). Exploring different mechanism in plant to resist drought and salt stress is essential to enhance crop resilience. One of the common plant responses to salt and drought stress is the production and accumulation of organic compounds called osmoprotectants or compatible solutes. These include proline (Pro), glycine betaine, choline, O-sulfate, sugars, and polyols (Silveira, Viégas Rde et al. 2003; Hoque, Okuma et al. 2007; Iqbal, Umar et al.2015). Proline is one of the important soluble osmotic regulator in plants under stressful conditions like drought (Urmi, Islam et al. 2023), salt (Huang, Bie et al. 2009), and heavy metals stress (Siripornadulsil, Traina et al. 2002). Some of proline’s unique functions arise from its inherent properties, while others depend on the coordinated “proline cycle” involving proline synthesis in chloroplasts and cytoplasm (Verslues et al. 2010).

Proline is considered a multifunctional amino acid, which is crucial for protein biosynthesis, serving as an osmotic protective compound, stabilizing macromolecules, acting as a free radical scavenger, and maintaining redox balance (Szabados, et al. Verslues, et al.2010). Various mutants or transgenic plants with proline deficiency display stunted growth, morphological abnormalities, altered flowering time, reduced fertility, or abnormal embryo development, suggesting that well-controlled proline metabolism is critical for vegetative development as well as for sexual reproduction (Mattioli, R., et al 2012Biancucci, et al.2015 Mattioli, R., et al. 2018). The accumulation of proline in plants depends on a balance between its synthesis and degradation processes (Szabados et al. 2010). The intracellular concentration of proline is regulated through metabolic synthesis (Roosens, Thu et al. 1998). Under stress condition in plants, there are two main pathways for proline synthesis (Hu, Delauney et al. 1992). Such as Proline is primarily produced through the glutamate pathway. In this process, glutamate is first phosphorylated and then reduced by pyrroline-5-carboxylate synthase (P5CS) to form pyrroline-5-carboxylate (P5C). P5C is then further reduced to proline by P5C reductase (P5Cr) (Armengaud, Thiery et al. 2004). Proline degradation starts with its oxidation by ProDH, which converts it to P5C. P5C is then further converted to glutamic acid by P5C dehydrogenase. Alternatively, proline can be synthesized via the ornithine pathway (Verslues et al. 2010). In this pathway, ornithine is first transformed into GSA and P5C by ornithine delta-aminotransferase (OAT), and P5C is then converted into proline (Roosens, Thu et al., 1998). ProDH enzyme responsible for the first step in proline degradation, plays a critical role in controlling proline accumulation and enhancing stress resistance in plants Two *ProDH1* and *ProDH2* genes was identified in *Arabidopsis harbor* (Funck, Eckard et al. 2010). Previous studies indicated that *ProDH1* is predominantly expressed in mitochondria, while *ProDH2* is expressed in both mitochondria and plastids (Kiyosue, Yoshiba et al. 1996, Van Aken, Zhang et al. 2009). The expression of the antisense *ProDH* gene in *Arabidopsis* was reduced, leading to an increase in proline accumulation (Nanjo, Kobayashi et al. 1999). ProDH and P5CR work together to mitigate the hypersensitive response in *Arabidopsis*, while also playing a key role in strengthening the plant’s defense mechanisms. We found that both *ProDH1* and *ProDH2* are necessary for maximum resistance against avrrpm1 and *Botrytis cinerea*. However, *ProDH2*2 plays a more significant role in early limiting the growth of *Metarhizium anisopliae* (Monteoliva, Rizzi et al. 2014).

The relative expression levels of proline catabolism genes (*ProDH1*, *ProDH2*) increased in sensitive varieties (add varieties name here). (Rizzi, Cecchini et al. 2016). The addition of proline and its analogue, thiazolidine-4-carboxylic acid (T4C), both substrates of ProDH, can further intensify and expose plant responses. Under drought stress, treatment with T4C in the drought-tolerant variety EC-98 leads to increase mRNA levels of *ProDH* genes, which are involved in both proline production and breakdown. The increase in *ProDH* gene expression may be linked to the conversion of T4C to cysteine, potentially contributing to the tolerance phenotype (Furlan, Bianucci et al. 2020;Dietrich, Weltmeier et al. 2011) found that *bZIP1* and *bZIP53* can bind to the promoter of the *ProDH* gene in *Arabidopsis*, playing a key role in reducing *ProDH* levels.

Similarly, (Dietmar Funck et al. 2010) observed that when *bZIP11* was inhibited by sucrose, the transcription of *AtProDH2* was also suppressed. *AtProDH2* is a target gene for *bZIP* in *Arabidopsis* (Hanseen et al. 2008). The interaction between the unknown gene *DFR1* and *AtProDH* in *Arabidopsis*, regulating drought stress. This suggests that under salt and drought stress conditions, *AtProDH* affects complex regulatory network mechanisms to cope with salt and drought stress.

Cotton is the most important fiber crop. Since it was first cultivated between 5000 and 10,000 years ago, cotton production has enormously influenced global economic development (Stephens SG et al. 1974). Proline catabolism mediated by Proline Dehydrogenases (GhProDHs) negatively impacts salt and drought tolerance in *Gossypium hirsutum* by accelerating proline degradation, thereby limiting the availability of proline for osmoprotection, reactive oxygen species (ROS) scavenging, and cellular homeostasis under abiotic stress. It is hypothesized that the downregulation or inhibition of *GhProDH* activity will enhance proline accumulation, improving the plant’s physiological response to salt and drought stress, resulting in better stress tolerance. Investigating the regulatory role of *GhProDHs* in proline catabolism will provide insights into how proline homeostasis can be manipulated to enhance cotton’s resilience to harsh environmental conditions. Therefore, the current study was aimed to: (1) Characterize the role of GhProDH2 in proline catabolism in *Gossypium hirsutum* under salt and drought stress conditions. (2) Determine the impact of *GhProDH2* gene expression on proline accumulation and stress tolerance in *Gossypium hirsutum.* (3) Explore the physiological effects of manipulating *GhProDH* activity on plant growth and survival under salt and drought stress. This research aims to unravel the complex role of proline catabolism in *Gossypium hirsutum* and provide a foundation for developing stress-tolerant cotton varieties through the manipulation of proline metabolism.

## 2. Results

### 2.1. Proline enhances cotton’s drought and salt tolerance

In order to investigate the effects of proline on cotton under salt and drought conditions, we used Proline and Proline inhibitors (ELB). We treated three leaf stage cotton seedlings with ddH_2_O, 200 mM NaCl, 15% PEG, Proline + 200 mM NaCl, Proline + 15% PEG, ELB + 200 mM NaCl, ELB + 15% PEG for 8 h. The results indicated that under ddH_2_O conditions, plants exhibited the best growth performance while under ELB + 200 mM NaCL condition, plants showed wilting on the leaf surface first, followed by leaf curling and showed wilting of all plant after 8 h. Under salinity stress, cotton plants exhibited moderate wilting. However, when salinity stress was combined with proline treatment, the wilting was less severe, indicating only mild wilting. Under ELB + 15% PEG treatment, cotton plants exhibited severe wilting while, moderate wilting was observed under PEG-only stress. When proline was applied alongside PEG, the cotton leaves showed less wilting, indicating that proline may help mitigate the negative effects of drought stress and improve the plant’s tolerance to such conditions.

Malondialdehyde (MDA) content increased under ELB + NaCl condition while decreased under proline treatment of cotton leaves (Figure 1). ELB Under drought stress, the MDA content in cotton was highest under the ELB + 15% PEG treatment, followed by the cotton leaves treated with 15% PEG. The MDA content in cotton was found to be lowest under the Proline + 15% PEG treatment, indicating reduced oxidative damage. Under salt and drought stress conditions, the activities of antioxidant enzymes such as SOD, POD, and CAT were highest when proline was added, followed by treatments with ELB.

**Figure 1.**
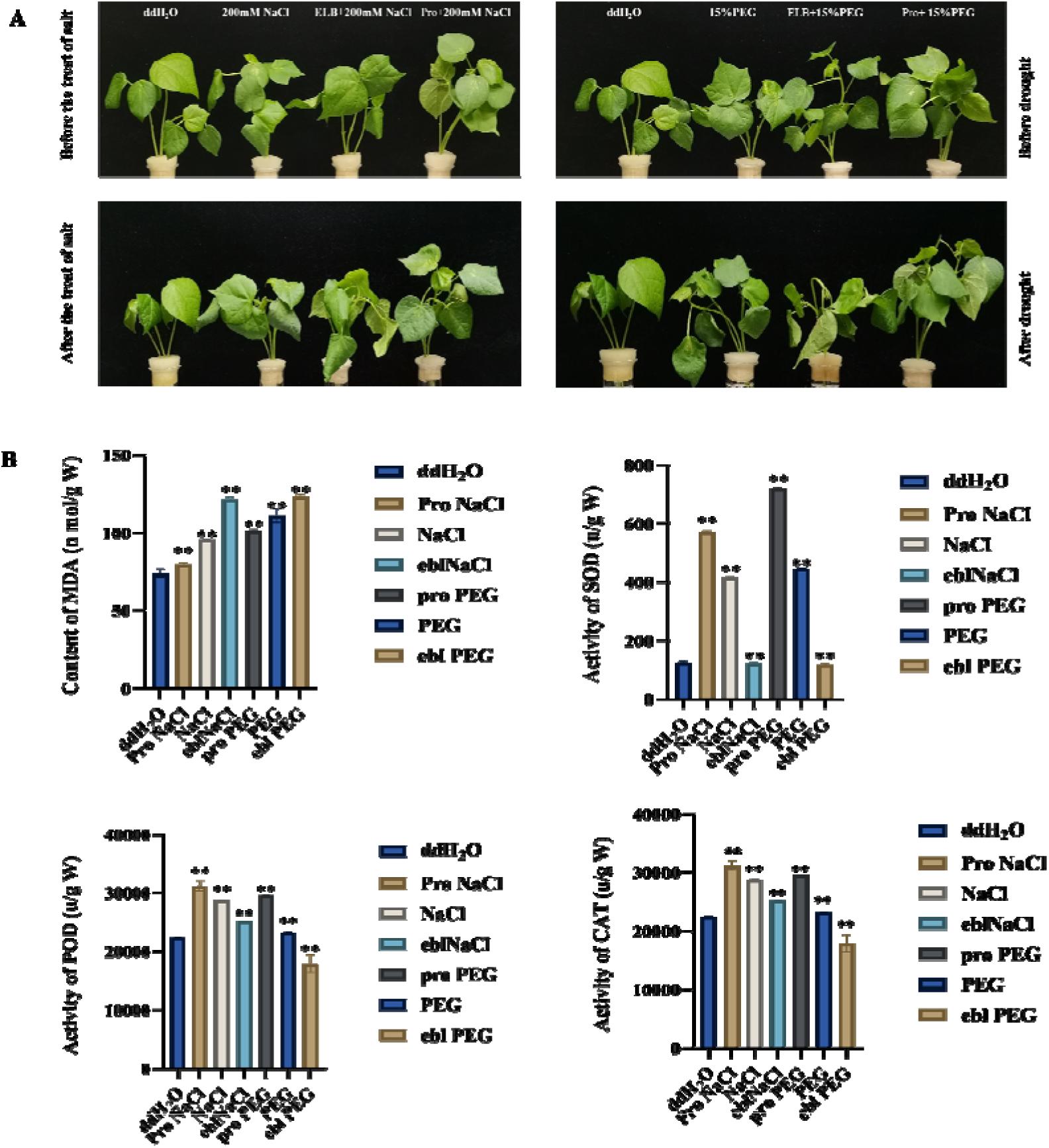
The effect of proline on drought and salt tolerance of cotton (A) Growth status of three leaf cotton under control conditions of ddH_2_O, salt stress (200mM NaCl), drought stress (15% PEG), and proline+salt stress (Proline+200mM NaCl), proline+ drought stress (Proline+15% PEG), Proline inhibitor+ salt stress (ELB+200mM NaCl), Proline inhibitor+ drought stress (ELB+15% PEG).(B) MDA content under various treatments,and SOD, POD, CAT content under various treatments Error bars represent ± SD of at least three biological replicates (* *P* < 0.05, ** *P*< 0.01, *** *P* < 0.001).”

### 2.2. Evolutionary trajectories、 collinearity analysis 、gene structure、 cis-Acting element and protein structure of ProDH in Cotton Species

The evolutionary relationship of *ProDH* genes in four cotton species showed a total of 18 *ProDH* genes.The analysis revealed that upland cotton and Sea island cotton each possess 6 *ProDH* genes, while Asian cotton and Raymond cotton have 3 *ProDH* genes each. The *ProDH* genes was clustered into three groups among four different cotton species. In upland cotton, *GhProDH2-1* was more closely to *GhProDH2-2*, *GhProDH2-3* to *GhProDH*, and *GhProDH2-5* to *GhProDH2-6.* According to collinearity analysis, *AtProDH* genes locate on chromosomes 3 and 5(Figure 2A). *GbProDH* and *GhProDH* genes are located on chromosomes A07, A08, A11, D07, D08 and D11, while *GaProDH* genes were found on chromosomes 07, 08, 11 and 01, 04, 07 respectively. Among *Arabidopsis* and the four cotton species has no collinearity relationship between *AtProDH1* and the cotton species. The *ProDH* gene in cotton aligns only with *AtProDH2*, demonstrating a linear correlation among *ProDH* genes in *G. hirsutum*, *G. arboreum*, *G. raimondii*, and *G. barbadense* (Figure 2C). Each *ProDH* gene in *G. arboreum* and *G. raimondii* corresponds to two *ProDH* genes in *G. hirsutum* and *G. barbadense*, respectively, further confirming that *ProDH* gene duplication occurred during the evolutionary process.The genome sequences and CDS sequences of *ProDH* genes in *Arabidopsis* thaliana, maize, rice, cotton, and Chinese cabbage,The gene structure was analyzed (Figure 2B). The vast majority of ProDH genes contain four exons, both Arabidopsis and cotton have four exons.

**Figure 2.**
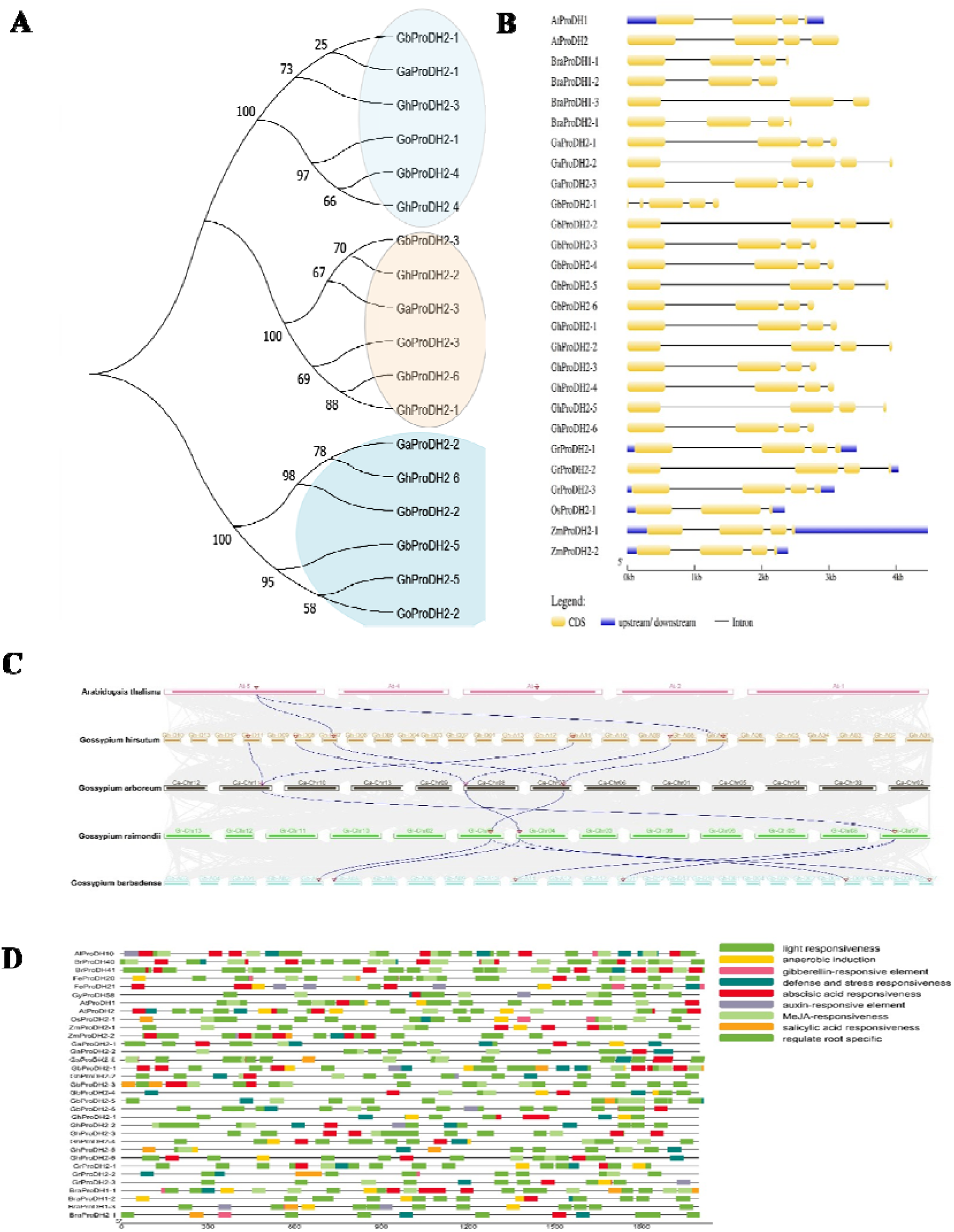
(A) Evolutionary relationships among four cotton species (B) The gene structure of *ProDH* genes in *Arabidopsis*, Maize, rice, cotton, and Chinese cabbage was analyzed (C) Collinearity analysis of *AtProDH*, *GhProDH*, *GaProDH*, *GrProDH*and *GoProDHgenes* (D) Analysis of Cis-regulatory element in *ProDH* genes.

Cis-acting elements are the binding sites of transcription factors, which regulate the precise initiation and efficiency of gene transcription. 2000 upstream sequence of *ProDH* genes were extracted to perform cis-acting element analysis. Cis-acting elements from 2000 upstream sequence of *ProDH* genes showed their diversified roles potential especially in various light responsiveness, anaerobic induction, gibberellin-responsive, abscisic acid responsiveness, defense and stress responses, MeJA-responsiveness, auxin-responsive element, salicylic acid responsiveness, regulate root specific nine categories (Fig. 2D). Notably, . All *ProDH* genes contain light response elements, and most of them contain stress related cis-elements except *ZmProDH 2-1*, *GbProDH 2-3*, *GhProDH 2-3* and *GrProDH 2-3*. This indicates that, *ProDH* genes are associated with stress response mechanism in plants.

We predicted the protein structure of ProDH with GMQE above 0.83 and sequence identity above 0.82, and analyzed six protein structures. We found that *GhProDH2-1* is structurally similar to *GhProDH2-2, GhProDH2-3* is structurally similar to *GhProDH2-4*, and *GhProDH2*-5 is structurally similar to *GhProDH2-6*(appendix Figure 1).

### 2.3. Expression Analysis of *GhProDH* Genes in Cotton

To understand the impact of *GhProDH2* on salt and drought tolerance in cotton, RNA seq data from the database (https://ngdc.cncb.ac.cn/) was analyzed which showed that *GhProDH2* were significantly expressed in tissues such as roots, stems, and leaves (appendix Figure 2A). Compared to other tissues, each *GhProDH2* gene exhibits higher expression levels in leaves, with *GhProDH2-1*, *GhProDH2-4*, and *GhProDH2-6* showing more expression . Expression patterns of *GhproDH* genes under salt and drought stresses at different time intervals (1h, 3h, 6h, 12h, and 24h) illustrates the negative trend of expression level with increasing time span, indicating their possible role in stress response (appendix Figure 2B). qRT-PCR analysis was conducted for confirmation of the RNA-seq results (appendix Figure 2C D). The expression level of *GhProDH2* genes under salt and drought stress was lower than control treatment. Due to the absence of stress related cis acting elements in *GhProDH2-3*, we selected five genes, *GhProDH2-1*, *GhProDH2-2*, *GhProDH2-4, GhProDH2-5,* and *GhProDH2-6* for further investigation.

### 2.4. Functional Analysis of *GhProDH* Genes in *Arabidopsis*

To investigate the biological functions of *GhProDHs*, we utilized *Agrobacterium* mediated transformation to produce overexpressed lines (OE) for *GhProDH2-1*, *GhProDH2-2*, *GhProDH2-4*, *GhProDH2-5* and *GhProDH2-6* as well as mutant lines for *AtProDH2-1*, *AtProDH2-2*, *AtProDH2-4*, *AtProDH2-5*, *AtProDH2-6* in *Arabidopsis* to explore their effects on salt and drought stress. The growth status of wilt-type (WT), OE (*GhProDH2-1*, *GhProDH2-2*, *GhProDH2-4*, *GhProDH2-5*, *GhProDH2-6*), and Mutant (*AtProDH2 -1*, *AtProDH2-2*, *AtProDH2-4*, *AtProDH2-5*, *AtProDH2-6*) lines were similar in 1/2MS medium. However, under 250mM Mannitol PEG and 125 mM NaCl stress, the growth status (Figures 3A), germination rate (Figure 3B), and root growth (Figures 3C and 3D) of overexpressed (*GhProDH2-1, GhProDH2-2, GhProDH2-4, GhProDH2-5, GhProDH2-6*) plants were worse than WT plants. Moreover, The overexpressed(*GhProDH2-1, GhProDH2-4, GhProDH2-5, GhProDH2-6*) plants were worse than The overexpressed(*GhProDH2-2*).The Mutant (*AtProDH2 -1, AtProDH2-2, AtProDH2-4, AtProDH2-5, AtProDH2-6*)plants were better than WT plants, Moreover, The Mutant (*AtProDH2 -1, AtProDH2-4, AtProDH2-5, AtProDH2-6*)plants were better than The Mutant(*AtProDH2-2*). So, the response of gene *GhProDH2-2* to salt and drought is not as strong as that of gene *GhProDH2-1, GhProDH2-4, GhProDH2-5, GhProDH2-6*.

**Figure 3.**
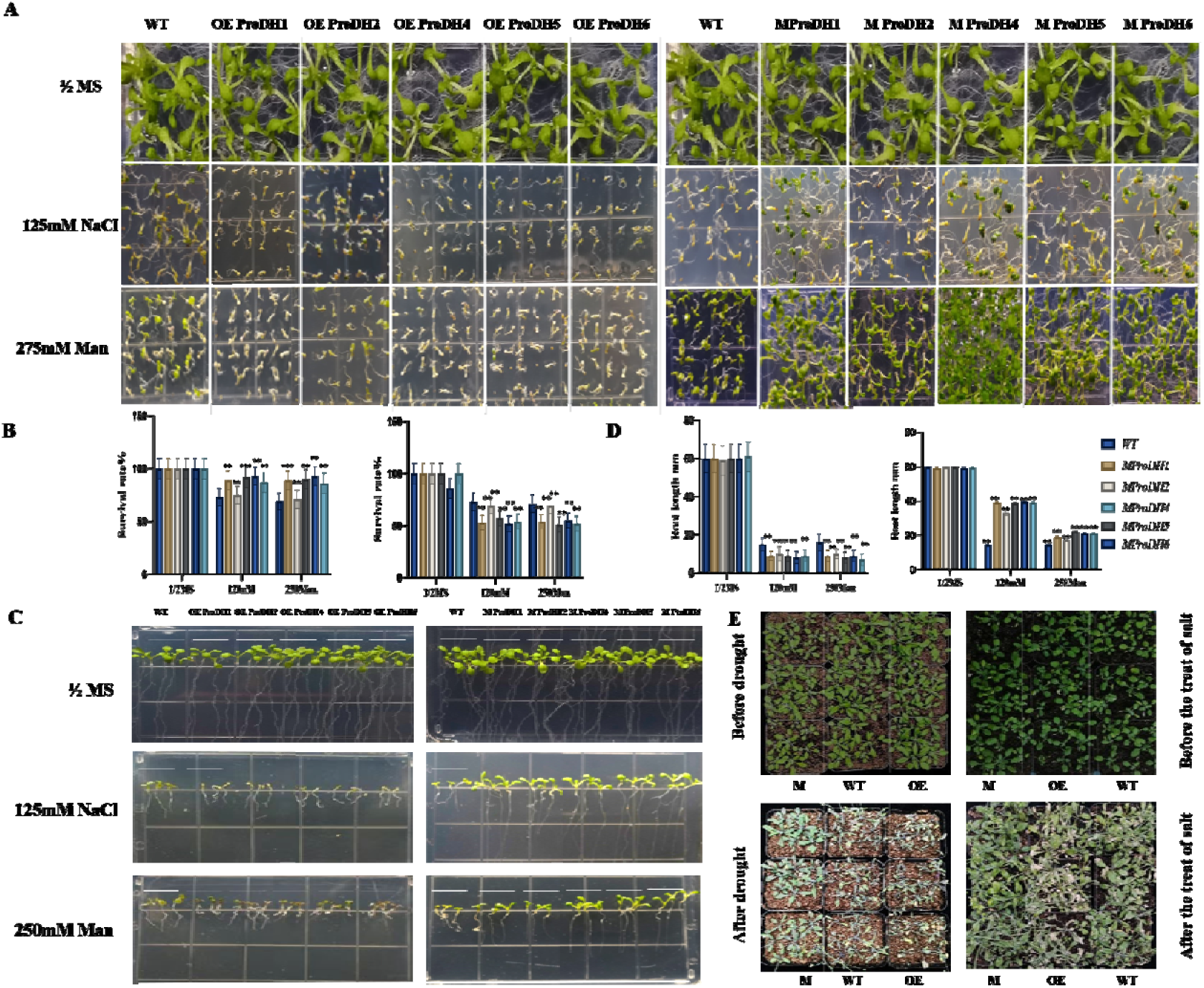
Functional validation of genetically modified *Arabidopsis*. (A) Germination assays were conducted using transgenic, mutant, and wildtype *Arabidopsis* seeds on ½ MS medium supplemented with 125 mM NaCl and 250 mM mannitol. The untreated ½ MS medium served as the control. (B) Survival rates were assessed for transgenic, mutant, and wildtype seeds exposed to 125 mM NaCl and 250 mM mannitol stress conditions. (C, D) Root length comparisons were made between transgenic, mutant, and wildtype plants grown on ½ MS medium supplemented with 250 mM mannitol and 125 mM NaCl for 10 days. (E) Response of *GhProDH2-4* transgenic lines and wildtype plants under drought and salt stress conditions. The error bars indicate ± SD of at least three biological replicates (* p < 0.05, ** p < 0.01, *** p < 0.001).

under 250 mM Mannitol PEG and 125 mM NaCl stress, the growth status (Figures 3A), germination rate (Figure 3B), and root growth (Figures 3C and 3D) of overexpressed lines (*GhProDH2-1, GhProDH2-2, GhProDH2-4, GhProDH2-5, GhProDH2-6*) plants were worse than WT plants. The Mutant (*AtProDH2 -1*, *AtProDH2-2*, *AtProDH2-4*, *AtProDH2-5*, and *AtProDH2-6*) plants were better than WT plants. In addition the response of gene *GhProDH2-2* to salt and drought is not as strong as that of gene *GhProDH2-1, GhProDH2-4, GhProDH2-5, GhProDH2-6*.

To further validate the results, *OE* and mutant *Arabidopsis* strains of *GhProDH2-4* were randomly selected and grown in medium for one week before being transferred to soil.

Without treatments, there was no significant differences occurred in the growth of WT, OE (*GhProDH2-4*), and mutant (*AtProDH2-4*) strains (Figure 3 E). To assess the role of *GhProDH4* in drought resistance, plants were grown in well-watered soil for 10 days before being subjected to drought conditions. After 10 days without water, the OE (*GhProDH2-4*) strain showed severe wilting, the WT strain showed moderate wilting, and the mutant (*AtProDH2-4*) strain showed mild wilting. Similarly, after 10 days of well-watered growth, no significant differences were observed in plant growth. However, following 15 days of salt stress with 125 mM NaCl, the OE strain exhibited severe leaf yellowing, the WT strain showed moderate yellowing, and the mutant strain showed only mild yellowing.

### 2.5. Functional Characterization of *GhProDH* Genes in Cotton

To investigate the function of *GhProDH2* in cotton, we employed virus-induced gene silencing (VIGS) to reduce the expression of the *GhProDH2* gene. Successful gene knockdown was confirmed by the bleaching of cotton plants infused with pYL156-PDS (Figure 4B). The expression levels of *GhProDH2-1*, *GhProDH2-4*, *GhProDH2-5*, and *GhProDH2-6* in the knockdown plants were significantly lower than those in the control plants containing the empty vector (pYL156) (Figure 4 C), indicating that all four genes were effectively silenced.

**Figure 4.**
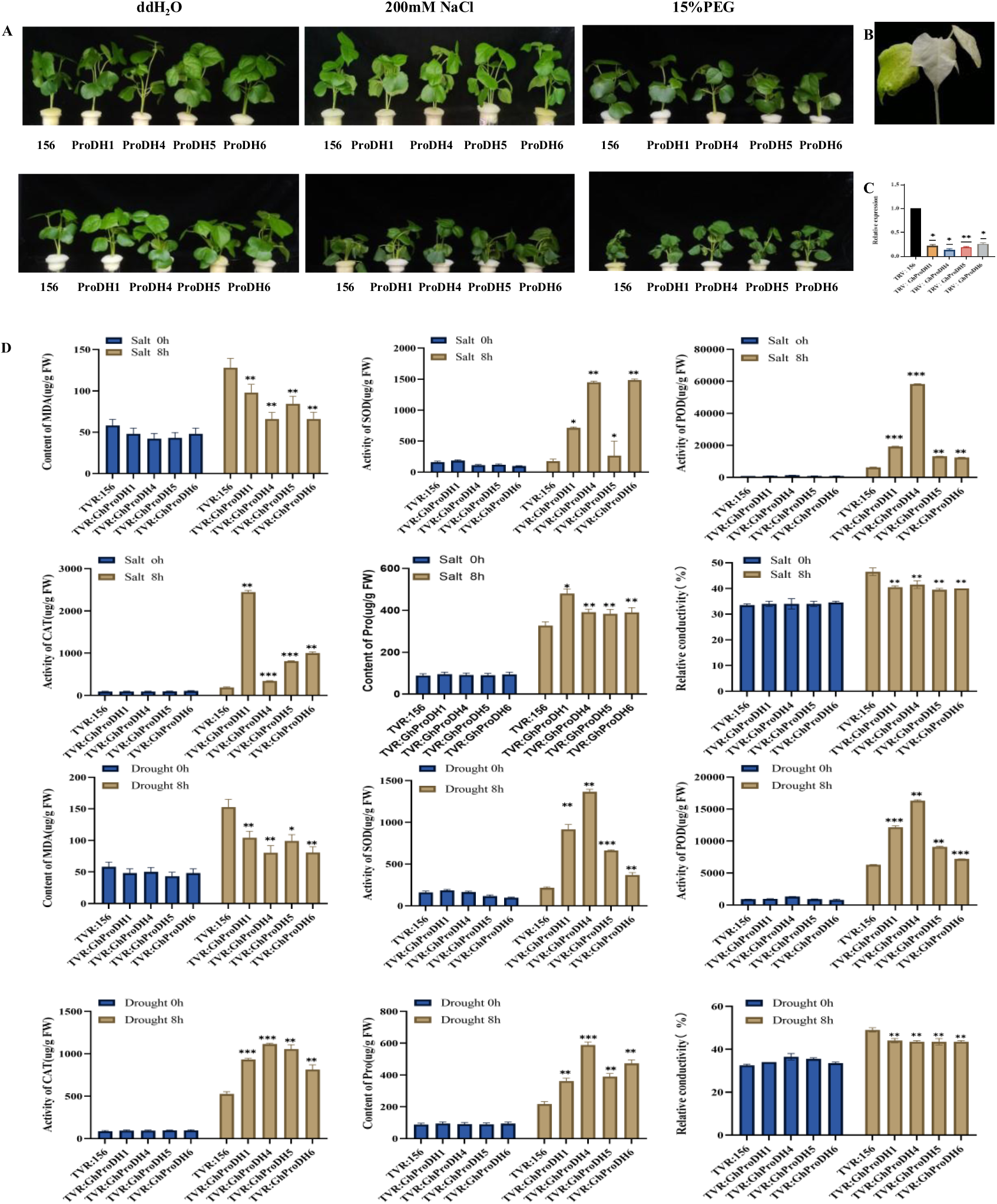
the impact of *GhProDH* silencing on cotton’s tolerance to drought and salt stress. (A)Comparision of phenotype of VIGS plants and control plants PYL156 before and after and control plants PYL156 before and after CK,200mM NaCl,15%PEG treatment. B) Representative albino marker plant used in the study. (C) Relative silencing efficiency assessed by qRT-PCR, showing VIGS plants with expression reduced to less than half compared to control plants. (D) Comparison of malondialdehyde (MDA) content between VIGS-silenced and control plants using pYL156 under drought or salt treatment. (E) Comparison of catalase (CAT), peroxidase (POD), superoxide dismutase (SOD), proline (Pro) contents and in VIGS-silenced plants versus control plants under drought or salt treatment conditions. Error bars represent ± SD of at least three biological replicates (* ***P*** < 0.05, ** ***P*** < 0.01, *** ***P*** < 0.001).”

Under 15% PEG treatment (simulated drought conditions), the control plants (pYL156) exhibited leaf curling earlier than other plants. Similarly, under 200 mM NaCl treatment (simulated salt stress), the control plants (pYL156) showed leaf wilting before other plants. (Figure 4A). Under salt stress, the wilting degree of GhProDH2-4 stems and leaves is milder than other genes (GhProDH2-1, GhProDH2-5, GhProDH2-6). Under drought stress, the curling degree of GhProDH2-4 leaves is lighter than other genes (GhProDH2-1, GhProDH2-5, GhProDH2-6). In addition, SOD, POD, Proline, CAT, relative conductivity and MDA content were measured in VIGS mediated (*pYL156 GhProDH2*) silent plants and control plants (pYL156). Under normal conditions, there was no significant difference in SOD, POD, Proline, relative conductivity and CAT content, but under salt and drought stress, the VIGS plants exhibited higher levels of these activity compared to the pYL156 control plants. This indicates that under salt and drought stress, VIGS silenced plants accumulate less ROS and increase cell osmotic pressure. Conversely, the VIGS plants accumulated less MDA content (Figure 5D). These findings indicate that the silencing of the *GhProDH2-1* 、 *GhProDH2-4* 、 *GhProDH2-5* 、 *GhProDH2-6* gene improve drought and salt stress resistance of cotton .

**Figure 5:**
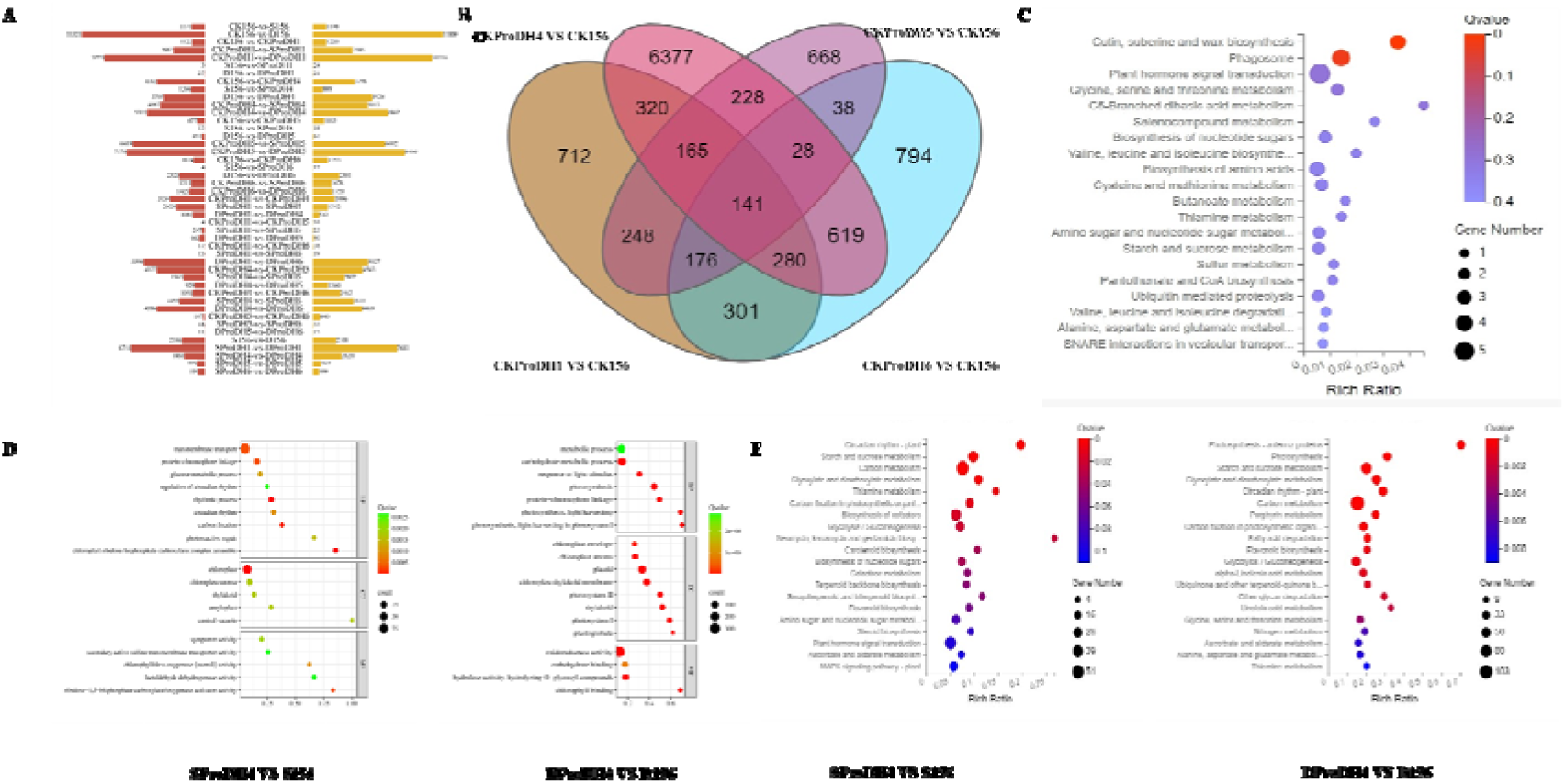
Detailed analysis of RNA-seq data depicting differentially expressed genes (DEGs) in various comparison sets. (A) Bar graph illustrating total DEGs, with red indicating upregulated and yellow-accented representing downregulated DEGs in each comparison. (B) Gene Venn diagram analysis under stress free conditions. (C) KEGG enrichment analysis of 141 genes. (D) GO enrichment analysis ofGhProDH2-4/156,under salt stress. (E) GO enrichment analysis of GhProDH2-4/156, under drought stress. (F) GhProDH5/156, and GhProDH6/156 under salt stress. (G) KEGG enrichment analysis of GhProDH2-4/156, under drought stress. CK156= pYL156; D156= pYL156 drought 8h; S156= pYL156 salt 8h; CKProDH1= pYL156:*GhProH1*; DProDH1= pYL156: *GhProDH1* drought 8h; SProDH1= pYL156: *GhProDH1* salt 8h; CKProDH4= pYL156:*GhProH4*; DProDH4= pYL156: *GhProDH4* drought 8h; SProDH4= pYL156: *GhProDH4* salt 8h; CKProDH5= pYL156:*GhProH5*; DProDH5= pYL156: *GhProDH5* drought 8h; SProDH5= pYL156: *GhProDH5* salt 8h; CKProDH6= pYL156:*GhProH6*; DProDH6= pYL156: *GhProDH6* drought 8h; SProDH6= pYL156: *GhProDH6* salt 8h

### 2.6. Transcriptome Analysis of the impact of silencing *GhProDH* Genes on Plant Stress Responses

To explore the regulatory mechanism of *GhProDH2*s in plants, we conducted a transcriptome analysis involving (control) TRV:156 and (VIGS) *TRV:GhProDH2-1 GhProDH2-4*, *GhProDH2-5* and *GhProDH2-6* under normal (0h), drought and salt stress (8 h). The transcriptome data revealed the detection of 62,179 genes, characterized by high-quality sequencing with Q20 and Q30 base proportions averaging 97.10% and 91.17%, respectively. Alignment rates showed robust comparisons of 95.22% to the genome and 79.60% to the gene set (supplementary table1). Furthermore, we summarized the differential gene expression levels across the samples, highlighting upregulated genes in red and downregulated genes in yellow (Figure 5A). 27 DEGs were found in S156 vs S*ProDH2-1*, 2154 DEGs were found in S156 vs S*ProDH2-4*, 31 DEGs were found in S156 vs S*ProDH2-5*, 21 DEGs were found in S156 vs S*ProDH2-6*, 48 DEGs were found in D156 vs D*ProDH2-1*, 9133 DEGs were found in D156 vs D*ProDH2-4*, 316 DEGs were found in D156 vs D*ProDH2-5*, 4701 DEGs were found in D156 vs D*ProDH2-6*, it was found that S156 vs S*ProDH2-4 and* D156 vs DProDH2-4 had the most differentially expressed genes. A total of 141 DEGs as common genes were found by Wayne diagram analysis that was performed for CK*ProDH2-1*/CK156, CK*ProDH2-4*/CK156, CK*ProDH2-5*/CK156, and CK*ProDH2-6*/CK156 (Figure 5B). KEGG enrichment analysis of these common DEGs showed that the most significant pathways were cutin, suberine and wax biosynthesis, phagosome, plant hormone signal transduction (Figure 5C), indicating that these DEGs related to common metabolic regulatory pathways. Overall, our findings highlight the complex role of *GhProDH2* genes in plant stress responses, as revealed by comprehensive transcriptomic analyses. These analyses provide high-quality data and valuable insights into gene regulation under different environmental conditions.

### 2.8. Gene Ontology revealed biological functions of *GhProDH2* Genes

In order to gain a deeper insights into the differential expression of genes, we conducted Gene Ontology (GO) analysis. Under salt stress conditions, GO analysis confirmed a diverse array of biological processes (BP) affected by TRV: 156-8 h/TRV: VIGS (*GhProDH2*) -8 h. This analysis categorized findings into cellular components (CC), molecular functions (MF), and BP. Notably, the top 20 enriched pathways, based on Q values, were plotted (appendix figure 3A,B).

Under salt stress, compared with the control group TRV: 156-8h, the *GhProDH2-1* silencing group significantly enriched the pathways including phosphorylation (BP), protein kinase activity (MF), sequence specific DNA binding (MF), and ATP binding (MF); The pathways significantly enriched in *GhProDH2-4* include chloroplast (CC), carbon fixation (BP), and ribose-1,5-bisphosphate carboxylase/oxygenase activator activity (MF). The pathways significantly enriched in *GhProDH2-5* include RNA polymerase II transcription factor complex (MF), fatty acid alpha hydroxylase activity (MF), L-leucine: 2-oxoglutarate aminotransferase activity (MF), and L-leucine transaminase activity (MF). The pathways significantly enriched in *GhProDH2-6* include DNA binding transcription factor activity (BP), iron ion binding (BP), and sterol metabolic process (BP).

Under drought stress, the top 20 pathways enriched in *GhProDH2-1* were ranked according to the rise and fall of Q values (Figure 5D), including glyoxylate cycle (BP), integral component of plasma membrane (CC), sucrose metabolic process (BP), and sulfate transport (BP). *GhProDH2-4* is enriched in significantly enriched pathways such as chloroplastic thylakoid membrane (CC), plastid (CC), photosynthesis (BP), and photosystem II (CC). *GhProDH2-5* was significantly enriched in the regulation of jasmonic acid mediated signaling pathway (BP), polyketide biosynthetic process (BP), and biosynthetic process (BP) pathways. *GhProDH2-6* was enriched in significantly enriched pathways such as photosystem II (CC), chloroplast thylakoid membrane (CC), chloroplast (CC), and photosystem I (CC). It can be seen that the pathways of gene enrichment under salt and drought conditions are different . And it can be summarized that under stress conditions, *GhProDH2-4* can activate more pathways.

### 2.9. KEGG analysis revealed Functional Enrichment pathways of G*hProDH2* Genes

KEGG analysis of *GhProDH2-1* under salt stress significantly enriched the Q value ≤ 0.05 and six pathways mainly enriched in Indole alkaloid biosynthesis Nitrogen metabolism, Betalain biosynthesis. Similarly, *GhProDH2-4* significantly enriched the Q value ≤ 0.05 in 20 pathways and it was mainly enriched in Circadian rhythm plant, Starch and sucrose metabolism, Glycoxylate and dicarboxylate metabolism. (appendix Figure3C,D) *GhProDH2-5* significantly enriched the Q value ≤ 0.05 in 4 pathways and it was mainly enriched in Glucosinolate biosynthesis Plant hormone signal transduction, Valine, leucine and isoleucine biosynthesis. Also, *GhProDH2-6* Q value ≤ 0.05 was significantly enriched in two pathways, mainly enriched in Ribosome, Brassinosteroid biosynthesis.

Under drought stress, KEGG enrichment mapping showed that *GhProDH2-1* significantly enriched the Q value ≤ 0.05 in 5 pathways and was mainly enriched in Glyoxylate and dicarboxylate metabolites, Sphingolipid metaboliscarbon metabolism. *GhProDH2-4* significantly enriched the Q value ≤ 0.05 in 20 pathways and it was mainly enriched in photosynthesis antenna proteins Starch and sucrose metabolism, Glyoxylate and dicarboxylate metabolism, and Circadian rhythm – plant. Under drought stress, (Figure 5E) *GhProDH2-5* significantly enriched 13 pathways, mainly in Flavonoid biosynthesis Cysteine and methionine metabolism, and Phenylpropanoid biosynthesis. *GhProDH2-6* significantly enriched 20 pathways, mainly enriched in photosynthesis Photosynthesis - antenna proteins, and Carbon metabolism. Therefore, GhProDH2-4 is significantly enriched in both GO and KEGG pathways under these conditions, suggesting it influences more pathways than the other genes. Consequently, *GhProDH2-4* was selected for further study.

### 2.10. Impact of *GhProDH2-4* Silencing on Starch and Sucrose Metabolism and Carbohydrate metabolism Pathways Under Salt and Drought Stress Conditions

Silencing *GhProDH* influences pathways involved in starch and sucrose metabolism, affecting genes such as alpha, alpha trehalose phase synthesis [UDP forming] 5] [2.4.1.15], sucrose synthase [2.4.1.13, UTP glucose 1-phase uridylltransferase [2.7.7.9] and granule bound start synthesis 1. [2.4.1.242], chloroplast/amyloplastic [3.2.1.68] glucose endo-1,3-beta-glucosidase 3 [3.2.1.39], endolucanase 17 like [3.2.1.14], and alpha glucosidase [12.4.1.1] under stress conditions as indicated by clustering heat map analysis. (appendix Figure 4A) Notably, *DProDH2-4/D156* and *D156/CK156* exhibit contrasting trends compared to *SProDH2-4/D156* and *S156/CK156*, with some genes showing downregulation such as alpha-amylase-like. ed alterations in Galactose metabolism, Neomycin, Kanamycin, and gentamicin biosynthesis, Biosynthesis of nucleotide sugars, Fructose and mannose metabolism, and Cyanamide metabolism, suggesting that *GhProDH2-4* silencing can impact various cellular sugar synthesis pathways, thereby enhancing plant drought and salt tolerance.Analysis of clustering heatmaps for *S156/SProDH2-4, SProDH2-4/S156, S156/CK156,* and *SProDH2-4/S156* after *GhProDH* silencing reveals contrasting trends in gene expression. Conversely, under salt stress, genes encoding components such as pfk (*phosphofructokinase,*, *glyceraldehyde-3-phosphate dehydrogenase,* ribulose bisphosphate carboxylase large chain, phosphoglycerate kinase,phosphoglycerate kinase, show upregulation. These findings underscore the intricate regulatory role of *GhProDH2-4* in modulating gene expression patterns critical for carbohydrate metabolism, which response to adverse conditions.

### 2.11. Metabolic Regulation by *GhProDH2-4*: Insights into Glyoxylate Dicarboxylic Acid Metabolism Under Salt and Drought Stresses

Upon silencing *GhProDH2-4*, analysis using clustering heatmaps revealed an inverse trend between *SproDH2-4/S156* and S156/CK156. Similarly, under drought conditions, clustering heatmaps *DProDH2-4/D156* and *D156/CK156* also exhibited opposite trends. *aconitate hydratase*、*ribulose bisphosphate carboxylase large chain*、*acetate/butyrate--CoA ligase AAE7 genes upregulation*.

These findings underscore the pivotal role of *GhProDH2-4* in regulating metabolic pathways essential for plant adaptation to salt and drought stresses, particularly through glyoxylate dicarboxylic acid metabolism and its interconnected metabolic pathways.

### 2.12. Metabolic Regulation by *GhProDH2-4*: Insights into Flavonoid biosynthesis Under Salt and Drought Stresses

Upon silencing *GhProDH2-4*, analysis using clustering heatmaps revealed an inverse trend between *SproDH2-4/S156* and S156/CK156.Similarly, under drought conditions, clustering heatmaps *DProDH2-4/D156* and *D156/CK156* also exhibited opposite trends. *chalcone synthase 1, flavonol synthase/flavanone 3-hydroxylase, caffeoyl-CoA O-methyltransferase etc* genes upregulation.

### 2.13. Physiological Responses of *GhProDH2-4* Suppression in Cotton Under Drought Stress

Under stress conditions, the palisade tissue in the leaves of both TRV:156 and VIGS plants exhibited a transition from long columnar to short cylindrical or spherical shapes, accompanied by a shift from tightly packed to loosely arranged structures, leading to decreased tissue compactness over time. However, the extent of these changes in the palisade tissue of VIGS plants was less pronounced compared to TRV:156 plants. This observation suggests that VIGS-mediated suppression of *GhProDH2-4* may confer a mitigating effect under stress conditions (Figure 6A).

**Figure 6.**
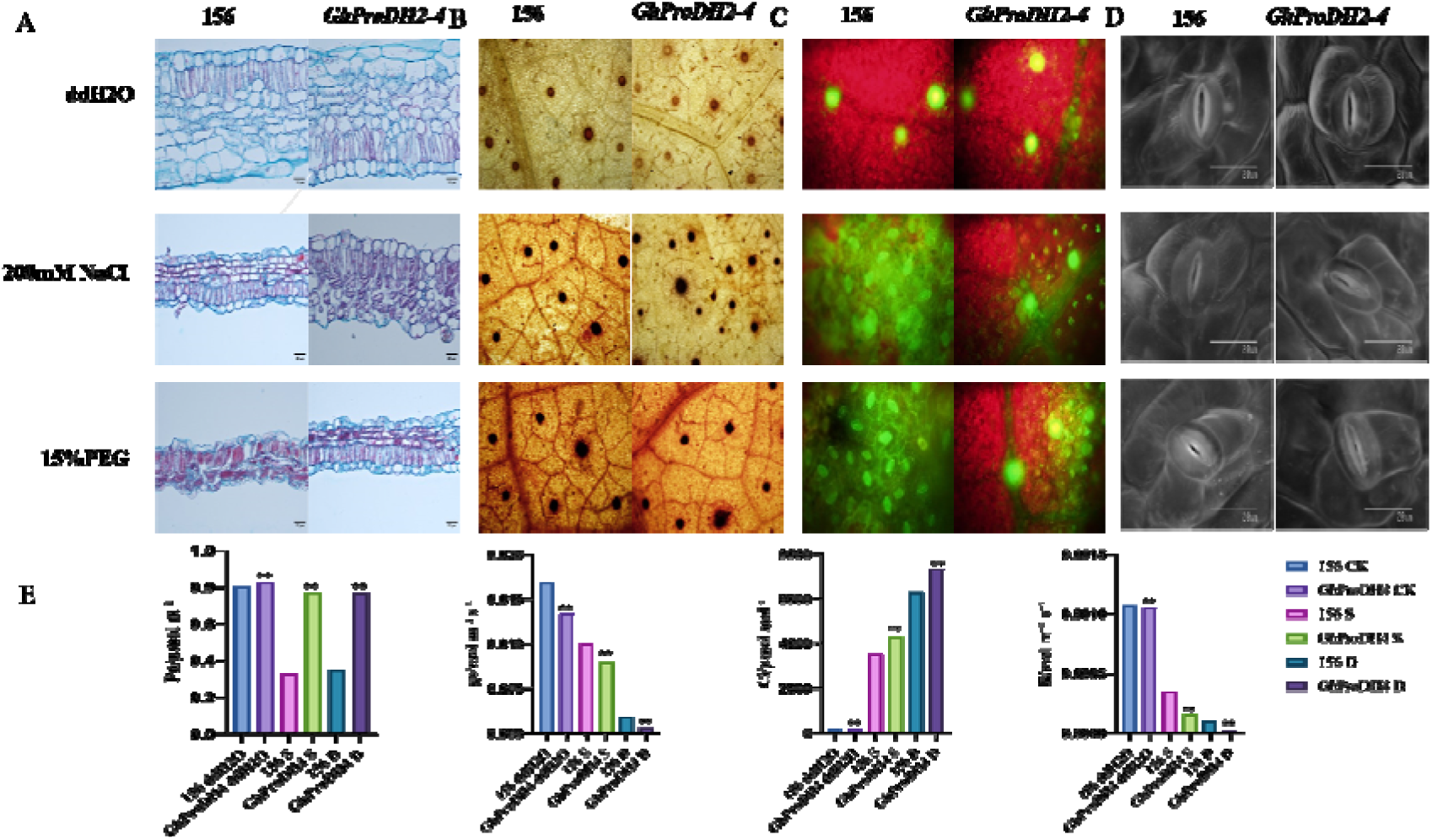
(A) Cotton plant TPV: 156, VIGS (GhProDH2-4) leaf cross-sectional condition under water, 200mM NaCl, and 15% PEG DAB staining method was used to detect the accumulation status of ROS in VIGS (GhProDH2-4) of cotton plants with TPV: 156 under water, 200mM NaCl, and 15% PEG conditions. (C) DCFH-DA method was used to detect the accumulation status of ROS in VIGS (GhProDH2-4) of cotton plants with TPV: 156 under water, 200mM NaCl, and 15% PEG conditions.(D)Cotton plant TPV: 156, stomatal status of VIGS (GhProDH2-4) under water, 200mM NaCl, and 15% PEG conditions. (E) Cotton plant TPV: 156, VIGS (GhProDH2-4) under water, 200mM NaCl, and 15% PEG conditions detect net photosynthetic rate (pn), intracellular carbon dioxide solubility (Ci), stomatal conductance (gs), transpiration rate (E).

ROS are inherently generated as byproducts of various metabolic processes such as photosynthesis and respiration. ROS levels are regulated by a network of enzymatic and non-enzymatic antioxidants distributed across cellular compartments. Additionally, ROS can traverse through aquaporins and other transport mechanisms to different cellular compartments or neighboring cells, where they function in signaling, detoxification, or accumulation processes (Mittler, Zandalinas et al., 2022). Given the enrichment of ROS-related processes in the extracellular domain under stress conditions, staining techniques were employed to assess ROS accumulation in TRV:156 and TRV plants. The results indicated that VIGS (*GhProDH2-4*) plants exhibited reduced ROS accumulation compared to *TRV:156* plants (Figure 6B). Further detection of intracellular ROS levels using the fluorescent probe DCFH-DA (Figure 6C) revealed that under ddH2O treatment, TRV: 156 and VIGS (*GhProDH2-4*) accumulated less fluorescence, while under salt and drought conditions, TRV: 156 accumulated more fluorescence than VIGS(*GhProDH2-4*).

Due to drought stress, *GhProDH2-4* is significantly enriched in pathways related to photosynthesis, where stomata serve as crucial regulators of plant gas exchange. Stomata can dynamically adjust their aperture in response to environmental changes, thereby influencing plant transpiration and photosynthetic activity (Wang, Li et al., 2024). It is hypothesized that *GhProDH2-4* may influence stomatal phenotype under drought conditions. Scanning electron microscopy analysis of both TRV:156 control plants and TRV (VIGS) plants revealed that under drought stress, the stomatal aperture of TRV plants was notably reduced compared to TRV:156 plants, indicating decreased stomatal opening in *GhProDH2-4*-suppressed plants (Figure 6D).

However, we measured the net photosynthetic rate, stomatal conductance, intracellular carbon dioxide concentration, and transpiration rate of 156 and VIGS plants under ddH2O, salt, and drought conditions using a photosynthesis analyzer 6800. We found that the stomatal conductance and transpiration rate of VIGS plants were lower than those of 156 plants, while the net photosynthetic rate and intracellular transpiration rate were higher than those of 156 plants (Figure 6E).

### 2.14. Metabolome analysis reveals the Regulatory Role of *GhProDH* Genes in Plant Stress Responses

To further study the metabolic changes in VIGS (*GhProDH2-4*) plants and TRV-156 plants under salt and drought conditions, 819 metabolites were identified in the metabolome with CAS numbers. These included 78 amino acids and derivatives, 130 phenolic acids, 100 flavonols, 54 flavonoids, 40 nucleotides and derivatives, 54 sugars and alcohols, 43 free fatty acids, 53 organic acids, and 53 alkaloids. (appendix figure 5). Under salt stress, KEGG analysis of TRV: 156-8 h/TRV: VIGS (*GhProDH*) -8 h revealed significant enrichment in Galactose metabolism ABC transporters, Aminobenzoate degradation, Flavonoid biosynthesis, carbon metabolism, Glyoxylate and dicarboxylate metabolism (Figure 7C). Under drought stress, KEGG analysis of TRV: 156-8 h/TRV: VIGS (*GhProDH2-4*) -8 h pathway revealed significant enrichment in carbon metabolism, flavonoid metabolism, and their compound protein digestion and absorption Galactose metabolism, Pyrimidine metabolism, carbon metabolism, Glyoxylate and dicarboxylate metabolism Pathways, these pathways include both primary and secondary metabolic pathways (Figure7D). Under salt and drought stress, the correlation between the enriched of different metabolites revealed a strong and extensive interactions between sugars, alcohols, flavonoids, organic acids, and other metabolites (Figure 7E-F). Primary metabolism can provide the necessary energy for the synthesis of secondary metabolites, such as under salt and drought conditions, which are significantly enriched in the carbon metabolism and glyoxylate dicarboxylic acid pathways. Thermal clustering analysis was conducted on flavonoids, sugars and alcohols, and organic acids, and it was found that the accumulation of these metabolites increased after gene silencing (Figure 7G).

**Figure 7.**
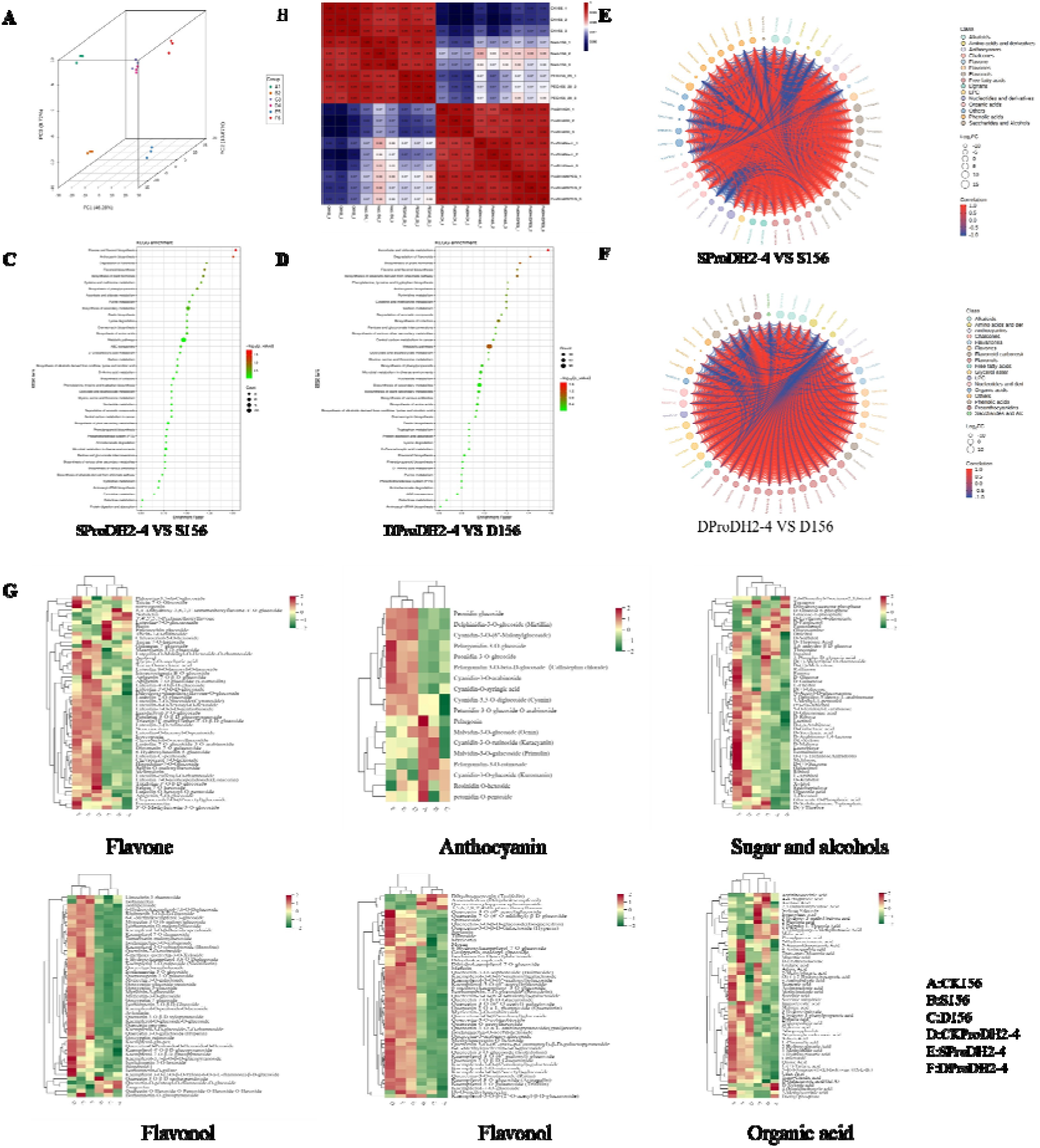
Principal component analysis (PCA) of metabolome profling of156CK,156NaCl,156 15%PEG,GhProDH2-4CK,GhProDH2-4NaCl, GhProDH2-415%PEG.B correlation of metabolome profling of156CK,156NaCl,156 15%PEG,GhProDH2-4CK,GhProDH2-4NaCl, GhProDH2-415%PEG (C) ProDH2-4 VS 156KEGG enrichment analysis under salt stress (D)ProDH2-4 VS 156KEGG enrichment analysis under drought stress(E) Correlation of differential metabolites under salt stress(F) Correlation of differential metabolites under drought stress(G) Cluster heatmap analysis of flavonoids, anthocyanin, flavonol, and organic acid metabolites in GhProDH2-4 and 156 plants under ddH_2_O, salt, and drought stress

### 2.15. Combined analysis of transcriptome and metabolome

Under salt and drought stress, silencing the *GhProDH2-4* gene significantly affected the carbon metabolism pathway, glyoxylate dicarboxylic acid metabolism pathway, flavonoid metabolism pathway, as well as the accumulation of sugars and alcohols in both transcriptome and metabolome. As a result, we concentrated on these three pathways and selected key genes and metabolites to compare their expression levels between VIGS (*GhProDH2-4*) plants and 156 plants under salt and drought stress conditions (Figure 8). We found that the metabolites of pyruvate and acetyl CoA, as well as soluble sugars increase in the carbon metabolism pathway. This can reduce the accumulation of oxidative active substances by altering cell osmotic pressure under salt and drought stress. In the flavonoid metabolism pathway, the accumulation of substances such as kaempferol, chalcone, anthocyanins, etc. can reduce the accumulation of ROS and enhance drought and salt tolerance. Organic acids such as malic acid, succinic acid, and glyoxylate accumulate in the dicarboxylic acid glyoxylate pathway.

**Figure 8.**
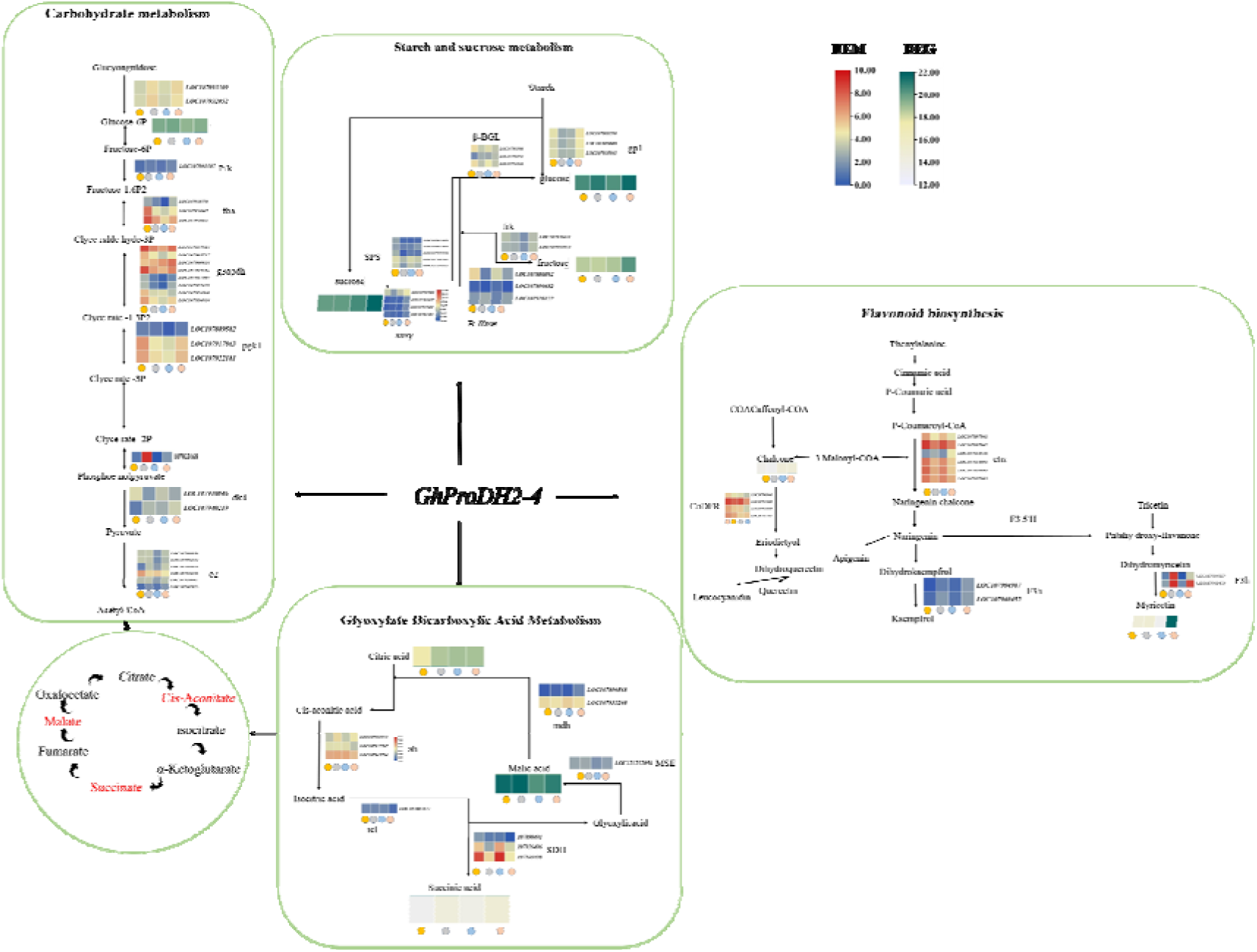
Changes in carbon metabolism, glyoxylate dicarboxylic acid, flavonoid metabolism pathway genes and metabolites. Note:pfk: Phosphpfructokinase; fba: fructose-bisphosphate aldolase, cytoplasmic isozyme 1;g3apdh: glyceraldehyde-3-phosphate dehydrogenase A; pgk1: phosphoglycerate kinase nse: pdk4: plastidial pyruvate kinase 4, e2: dihydrolipoyl dehydrogenase 2; mdh: malate dehydrogenase MSE: malate synthase, glyoxysomal-like SDH: succinate dehydrogenase icl: isocitrate lyase-likef3h: flavonol synthase /flavanone 3-hydroxylasechs: chalcone synthase 1; gp1: alpha-glucan phosphorylase 1;β-BGL: beta-glucosidase BoGH3B;frk: fructokinase-6;SPS: sucrose-phosphate synthase 4susy: sucrose synthase-like; B-ffase: acid beta-fructofuranosidase

## 3. DISCUSSION

Cotton is one of the most important fiber crops, accounting for 35% of global fiber (Abdelraheem et al. 2019). Drought alone is currently reported to affect about 45% of agricultural land worldwide; likewise, about 19.5% of the cultivable agricultural lands are under salinity stress (Dos Reis et al. 2012) which have a significant impact on the yield of cotton. Proline is an amino acid which in primary metabolism in plants, serves as one of the most widely distributed and effective organic compatible osmolytes (Agami et al. 2016; Mohamed et al. 2020). It accumulates in plant cells under stress conditions and plays an important role in plant stress resistance (Ben Rejeb et al.2012). Role of proline in enhancing the salt and drought tolerance in plants has been studied as it can participate in regulation of cell division, cell elongation, cell wall self-assembly to sustain a fast elongation for the hairy roots and other organs including reproductive system(Per, Tasir S. et al.2017; Singh et al.2015Duan et al. 2024; Trovato et al. 2008) . The growth performance of cotton plants was investigated with exogenous application of proline and ELB (proline inhibitor) under salt and drought stress conditions. Under salt and drought conditions, the growth performance of cotton plants was better under exogenous proline treatment than that under exogenous treatment of proline inhibitor(Figure 1.)The use of proline inhibitor (EBL) to treat transgenic lemons resulted in a decrease in salt tolerance, while the supply of exogenous proline partially restored the salt tolerance of the VIGS strain (Dai, W.et al.2018).ProDH, the rate-limiting enzyme in the first step of proline degradation pathway, has a certain impact on proline content. To investigate the regulatory mechanism of ProDH on salt and drought tolerance in cotton plants, we studied the function of *GhProDH2* gene. This study identified 18 *ProDH* genes from four cotton species and distributed them into three groups. Structural analyses of the *ProDH* gene revealed the similarities between motif and structure of *ProDH* genes. Specifically, the six *GhProDH2* gene structures in upland cotton contain four exons (Figure 2A).

A collinearity analysis was performed between *Arabidopsis* and four cotton species, revealing that the *ProDH* gene in cotton does not have a homologous pair with *AtProDH1*, but shows a certain linear correlation with *AtProDH2* (Figure 2C). This suggests a possible divergence in the evolutionary paths of these genes between cotton and *Arabidopsis*. To further understand the regulatory mechanisms of *ProDH* genes, a cis-acting element analysis was conducted on the 2000 upstream sequences of *ProDH* genes from various developmental stages of plants. The results identified nine categories of cis-acting elements: photo reactivity, anaerobic inducibility, gibberellin responsive elements, abscisic acid responsiveness, defense and stress responsiveness, MeJA responsiveness, auxin responsive elements, salicylic acid responsiveness, and regulatory root specificity. However, *GhProDH2-3* was found to have no stress cis acting elements in upland cotton (Figure 2D). This comprehensive analysis highlights the conserved structural features of *ProDH* genes in upland cotton and their potential regulatory elements, providing insights into their roles in stress responses and primary metabolism. Through protein structure analysis, it was found that *GhProDH2-1* is structurally similar to *GhProDH2-2*, while *GhProDH2-3* is structurally similar to *GhProDH2-4*, and *GhProDH2-5* is structurally similar to *GhProDH2-6* (Attach Figure 1). Transcriptome data showed that *GhProDH2-1*, *GhProDH2-2*, *GhProDH2-3*, *GhProDH2-4*, *GhProDH2-5*, and *GhProDH2-6* were all expressed in the plant leaves.

Through overexpression and silencing of *GhProDH2-1*, *GhProDH2-2*, *GhProDH2-4*, *GhProDH2-5*, and *GhProDH2-6*, The *GhProDH2* gene was found to have a negative regulatory effect on salt and drought tolerance. Due to the weaker response of *GhProDH2-2* to salt and drought stress compared to other genes. We next studied the drought and salt tolerance of *GhProDH2-1*, *GhProDH2-4, GhProDH2-5*, and *GhProDH2-6* in cotton. GhProDH2-4 has been found to have a strong response to salt and drought than other genes (*GhProDH2-1*, *GhProDH2-5*, and *GhProDH2-6*) Salt and drought stress generate higher ROS molecules which the antioxidant enzymes can effectively eliminate its production by metabolic activities(Meng, Y.C. et al. 2021) ROS (Uzilday, B. et al.) . The activity levels of superoxide dismutase, peroxidase, and catalase were higher in the silenced VIGS *GhProDH2-1*, *GhProDH2-4*, *GhProDH2-5*, and *GhProDH2-6* of cotton than that in the control group TRV: 156. These results are in agreement with several previous studies (Schekaleva, O. et al. 2024 ;Liu, C. et al. 2023). MDA, as one of the end products of lipid peroxidation reaction in organisms, is an important parameter reflecting the potential antioxidant capacity of the plant cells and can indirectly reflect the degree of tissue peroxidation damage (Xu, Y. et al. 2020; Tsikas, D. et al.2017). Transcriptome analysis was conducted on silenced plants (VIGS *GhProDH2-1, GhProDH2-4, GhProDH2-5, GhProDH2-6*) and the control (TRV: 156) in upland cotton. Under stress-free conditions, all silenced plants exhibited enrichment in cellular components and shared the same GO pathway, confirming their homology. However, each gene also displayed unique enriched pathways, highlighting their distinct and synergistic roles. GO and KEGG enrichment analyses under salt and drought stress revealed that VIGS (*GhProDH2-1, GhProDH2-4, GhProDH2-5*, and *GhProDH2-6*) plants had significantly enriched pathways compared to TRV: 156. Despite *GhProDH2-5* and *GhProDH2-6* being more closely related, their significantly enriched pathways were not more similar to each other than to those of the other genes. This indicates a complex interaction and distinct functional contributions of each *GhProDH* gene under stress conditions. The various genes of the GhProDH family (*GhProDH2-1, GhProDH2-4, GhProDH2-5,* and *GhProDH2-6*) regulate different pathways to collectively combat salt and drought stress. Notably, GhProDH2-4 is significantly enriched in both GO and KEGG pathways under these conditions, suggesting it influences more pathways than the other genes. Consequently, *GhProDH2-4* was selected for further study. It was found that silencing *GhProDH2-4* significantly affects the starch sucrose pathway, carbon metabolism pathway, glyoxylate dicarboxylic acid pathway, and flavonoid metabolism pathway under salt and drought stress.

Anatomical observations of cotton leaves under salt and drought conditions revealed that silencing GhProDH2-4 mitigates the tightening of sponge and palisade tissues (Figure 7). The *GhProDH2-4* gene is enriched in the photosynthesis pathway, where stomatal movement is essential for regulating photosynthesis rate, water balance, and plant immunity (Rodrigues et al. 2022). This research indicates that *GhProDH2-4* gene enhances resistance to salt and drought stress by modulating stomatal opening (Figure 7). ROS can damage proteins, lipids, and nucleic acids and can alter protein properties by forming covalent bonds, thus regulating their activity and function (Møller, Jensen et al. 2007). The production of ROS is a direct defense mechanism against pathogens, as they are toxic to pathogens and signal responses to biotic and abiotic stress stimuli or developmental cues (Wrzaczek, Brosché et al. 2013, Baxter, Mittler et al. 2014). Net photosynthetic rate can be used to characterize the health status of plants and their ability to accumulate organic matter (Zhang, Sun et al. 2022). Silencing *GhProDH2-4* reduces ROS accumulation, thereby enhancing resistance to salt and drought stress. Transcriptome analysis of silenced *GhProDH2-4* under stress conditions showed significant enrichment in starch and sucrose metabolism, carbohydrate metabolism, glyoxylate, and dicarboxylate metabolism pathways, flavonoid biosynthesis indicating that salt and drought stress share common signaling pathways.

Metabolomics study revealed that under salt and drought stress, VIGS (*GhProDH2-4*) was significantly enriched in carbon metabolism, flavonoid metabolism, and glyoxylate dicarboxylic acid pathway compared to TRV: 156. Flavonoids are essential antioxidants for plant growth, development, and reproduction, and can enhance plant stress resistance (Zhang, Huang et al. 2022; W. Yang Yang, et al. 2021). Carbon metabolism is not only crucial for plant growth and development, but also plays a key regulatory role in stress response (Li, Li et al. 2024;Fang, S. *et al*. 2023). Glyoxylate dicarboxylic acid metabolism plays a critical role in carbohydrate synthesis, serving as both an energy source and signaling molecules in response to abiotic stresses in plants (Zeb, Liu et al. 2022;Lv, H. *et al*. 2024). The enrichment of glyoxylate dicarboxylic acid metabolism is advantageous for accumulating osmotic substances, initiating osmotic regulation, and clearing ROS (Huang, Peng et al. 2021). It was found that there is a certain accumulation of metabolic sugars, alcohols, acetone metabolites, and organic acids in the metabolome. Through joint analysis of transcriptome, metabolome, and physiological phenotype, it was found that silencing the *GhProDH2-4* gene upregulates carbon metabolism pathway related genes to provide plants with certain energy, increase net photosynthetic rate, and accumulate soluble sugars. Increased concentrations of soluble sugars have been linked to enhanced drought (O’Brien, Valtat et al. 2020) and salt tolerance (Li, Meng et al. 2022) in plants, through controlling cell osmotic pressure. Silencing of *GhProDH2-4* gene also increases proline content, cell osmotic pressure and affects the glyoxylate dicarboxylic acid pathway, leading to the accumulation of organic acids and reducing ROS accumulation (Figure 9). Moreover, silencing the GhProD*H2-4* gene increases gene expression in acetone related pathways, leading to the accumulation of secondary metabolites such as flavonoids which depricing ROS accumulation, thereby jointly regulating cotton’s drought and salt tolerance (Figure 9).

**Figure 9.**
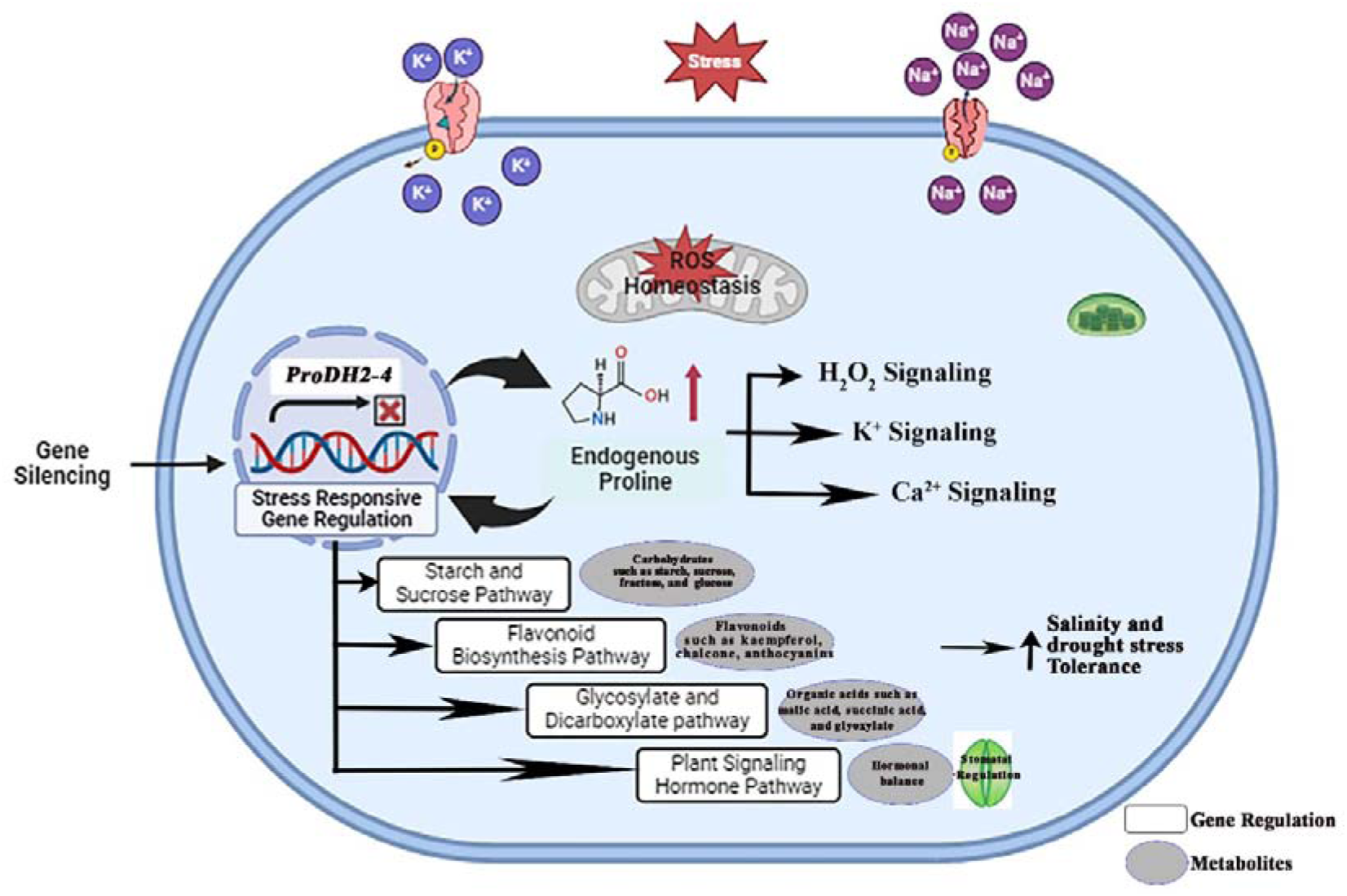
Regulatory network model of ProDH2-4 role in cotton plant under drought stress and salt stress.

## 4. CONCLUSION

This study explores how proline enhances the drought and salt tolerance of cotton. The ProDH gene family, as the rate limiting enzyme for proline degradation, has a certain impact on plant drought and salt tolerance. There was only *GhProDH2* gene in cotton, which has a certain degree of conservation. Evolutionary tree and protein structure prediction revealed that *GhProDH2-1, GhProDH2-3* and *GhProDH2-5* were more closely related to GhProDH2-2, GhProDH2-4, and *GhProDH2-6*, respectively. Overexpression and silencing of *GhProDH2-1, GhProDH2-2, GhProDH2-4, GhProDH2-5,* and *GhProDH2-6* genes in *Arabidopsis thaliana* tolerate the toxicity of drought and salt stress. In addition, the response of GhProDH2-2 gene to salt and drought stress is weaker than other genes(*GhProDH2-1, GhProDH2-4, GhProDH2-5,* and *GhProDH2-6)*. In cotton, VIGS silencing of *GhProDH2-1, GhProDH2-4, GhProDH2-5,* and *GhProDH2-6* genes enhanced cotton drought and salt tolerance. Under salt and drought stress,The wilting status of cotton leaves after silencing *GhProDH2-4* is milder than other genes(*GhProDH2-1, GhProDH2-5,* and *GhProDH2-6*) Transcriptome analysis revealed that under salt and drought conditions, *GhProDH2-4* had the most significant impact on DEGs. Through microstructure analysis, it was found that silencing *GhProDH2-4* alleviates leaf wrinkling, reduces ROS accumulation, decreases stomatal opening, and enhances net photosynthetic rate. Through transcriptome, metabolome and physiological phenotype joint analyses, it was found that silencing of *GhProDH2-4* enhances cotton’s drought and salt tolerance by affecting carbon metabolism, flavonoid metabolism, glyoxylate dicarboxylic acid pathway accumulation of soluble sugars, flavonoid metabolites, organic acids, and other substances. This study provides a theoretical basis for exploring role of ProDH in the drought and salt tolerance of plants. At the same time. provides valuable genetic targets for engineering crop varieties with highly stress resistant.

## 5. MATERIALS AND METHODS

### 5.1. Research design and treatments

To promote seed germination in the upland cotton variety “TM-1”, seeds were surface sterilized by using 95% concentrated sulfuric acid, and dried. The seeds were transplanted into pots containing a 2:1 mixture of nutrient soil and vermiculite. The growth conditions included a temperature of 25°C, humidity of 37%, a light cycle of 16/8 h, and a light intensity of 2000 lx in a constant temperature incubator. Seedlings were subjected to drought and salt treatments by using 15% PEG and 15% NaCl, respectively. Samples were collected at 1h, 3h, 6h, 12h, and 24h intervals, immediately frozen in liquid nitrogen, and stored at -80°C for RNA extraction.

For *Arabidopsis thaliana*, seeds were sterilized with mercury chloride and placed on 1/2 MS culture medium. They were kept in the dark at 4°C for 24 h, then transferred to a constant temperature incubator for about a week. Once the seedlings were growing well, they were moved to pots filled with a 2:1 mixture of nutrient soil and vermiculite. After 3 to 5 days of drought treatment, physiological indicators were measured. Seven days after treatment, the plants’ morphology was observed, and survival rates were recorded after 3 days of rehydration..

### 5.2. RNA extraction and qRT PCR analysis

To extract RNA from the prepared samples, the samples were grinded in liquid nitrogen and use the RNAprep Pure Plant Plus Kit. Measure the RNA concentration, and then use the TransScript One-Step gDNA Removal and cDNA Synthesis SuperMix kit to reverse transcribe the RNA into cDNA.Dilute the cDNA five times and use it as a template for fluorescence quantitative PCR (q-PCR). The reaction system and program details are provided in Tables 1-1 and 1-2. q-PCR primers are designed using the Real-Time PCR (TaqMan) Primer and Probes Design Tool website, with Ghactin serving as the internal reference gene. The relative gene expression levels are calculated using the 2-ΔΔCt method by three replicates.

### 5.3. Cloning of *GhProDH* gene

Coding sequence (CDS) were used for clone the *GhProDH* gene with specific primers(appendix table S2) . Using cDNA as the template for amplification the complete CDS sequence. Run the PCR product on a 1% agarose gel electrophoresis to obtain the electrophoresis product. Use the Megan Agarose Gel Recovery Test Kit to cut and recover the bands of the target fragment size, then purify them to obtain the target gene. Using the ClonExpress II One-Step Cloning Kit, ligate the target gene into the T vector. Transform the Vazyme #C502 competent cells through heat shock. Plate the transformed Escherichia coli onto LB medium containing kanamycin and incubate overnight at 37°C in an inverted position

### 5.4. Construction of VIGS and Overexpression Vectors for *GhProDH* Gene

Specific primers were used for overexpression and VIGS of the *GhProDH* gene. Using the extracted RNA Reverse transcript into cDNA as a template, the gene fragments were amplified and purified, then stored at -20 °C for future use. The overexpression vector PBI121 was cleaved at Hind III and BamHI sites, while the VIGS vector Pyl156 utilized BamHI and Sacl sites. Following gel electrophoresis and recovery, the linearized vectors underwent Fusion Ligation technology to form complete constructs. Subsequently, these vectors were transformed into *Escherichia coli*, where single clones were selected and sequenced for verification. Plasmids from confirmed sequences were extracted and transformed into Agrobacterium tumefaciens. After cultivation on LB agar supplemented with kanamycin and rifampicin, monoclonal colonies were selected and re-validated to ensure correct insertion of the *GhProDH* gene. Finally, the validated bacteria were preserved at -80°C for future experiments.

### 5.5. Agrobacterium-Mediated Transformation and Infiltration Process

The recombinant plasmids were transformed into Agrobacterium using the GV3101 Chemical Competent Cell reagent kit. Cultures of PYL156 PDS (positive control), PYL192 (auxiliary vector), PYL156 *GhProDH2-1* (positive control), PYL156 *GhProDH2-4* (positive control), PYL156 *GhProDH2-5* (positive control), PYL156 *GhProDH2-6* (positive control), and PYL156 (empty vector) were grown in liquid LB medium supplemented with rifampicin and kanamycin at 28°C with agitation at 200 RPM. They were shaken overnight until reaching an OD600 value of 1.0-1.5, then centrifuged and resuspended in a heavy suspension solution (MES, magnesium chloride, AS, water) at 25°C in darkness for 3-4 h. The PYL156 (EV), PYL156 *GhProDH2-1* (VIGS), PYL156 *GhProDH2-4* (VIGS), PYL156 *GhProDH2-5* (VIGS), PYL156 *GhProDH2-6* (VIGS), and PYL PDS suspensions were mixed with PYL192 in a 1:1 ratio. Each cotyledon’s underside was injected with the mixed solution, followed by incubation in a 25°C incubator in darkness for 12 h, then transferred to a light cycle of 16 h light/8 h dark.

### 5.6. Infection of Arabidopsis and Screening of Transgenic Arabidopsis

Agrobacterium tumefaciens carrying overexpression vectors was added to LB liquid medium containing kanamycin and rifampicin, and shaken at 28°C and 200 rpm until reaching an OD value of 1.0-1.5. After centrifugation, the bacterial pellet was resuspended in an equal volume of resuspension buffer (1/2MS, 5% sucrose, pH 5.8), adjusting the OD value of the suspension to 0.8. The suspension was treated in darkness at 25°C for 2 hours. Then, 0.02% Silwet L-77 was added to the suspension for infection. Flower buds were immersed in the suspension for about 1 minute, laid flat, and incubated in darkness for 24 hours before returning to normal growth conditions. This infection process was repeated after one week.

Following infection, seeds of *Arabidopsis thaliana* were designated as the T0 generation. These seeds underwent a 5-minute wash in mercuric chloride, followed by 7-8 washes in sterile water. They were then plated on 1/2MS medium containing kanamycin for 24 h of vernalization. Positive seedlings were selected and transplanted into small pots. The seeds obtained from these plants were labeled as the T1 generation, and the process was repeated until obtaining seeds from the T3 generation.

## Acknowledgements

This work was supported by the National Key R&D Program of China (2023YFE0102200), China Agriculture Research System of MOF and MARA(CARS-15-38), Biological Breeding-Major Projects in National Science and Technology (2023ZD04038), Key R&D Program of Shandong Province (2023LZGC007) and the National Natural Science Foundation of Xinjiang (2022D01A159).

## Author Contributions

Suli Bai: performed research, tools, analyzed data, wrote the paper and editing. Suli Bai and Pei Zhao: resources, investigation. Fei Su,Shuai Wang, Junjuan Wang and Yan Li: tools, visualization, computational. Ibrahim A. A. Mohamed and Zujun Yin: designed the research, contributed new analytic, writing-reviewing and editing. All authors have read and approved the final manuscript.

---

**appendix figure 1:**
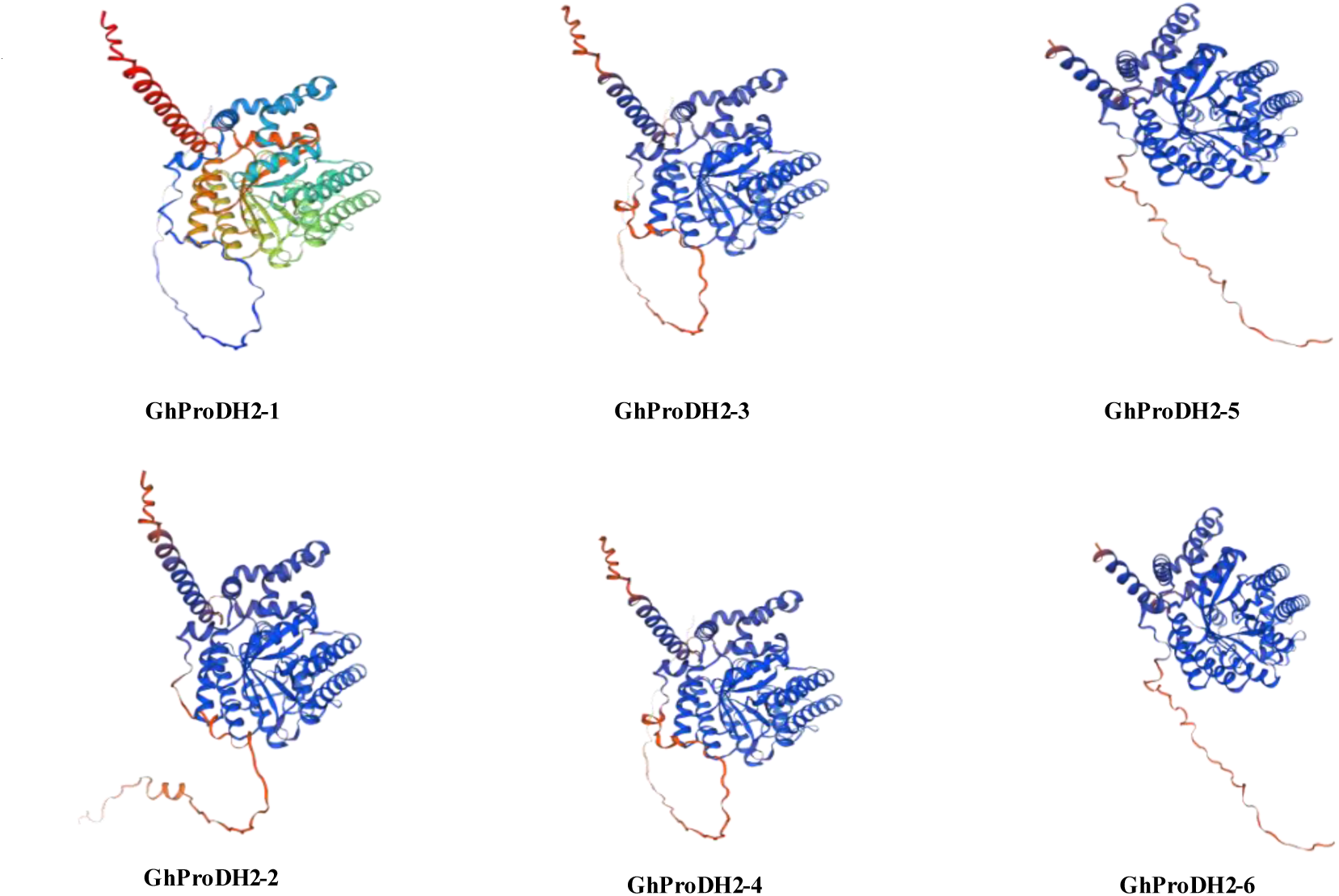
Prediction of *GhProDH* protein structure in upland cotton.

**appendix figure 2.**
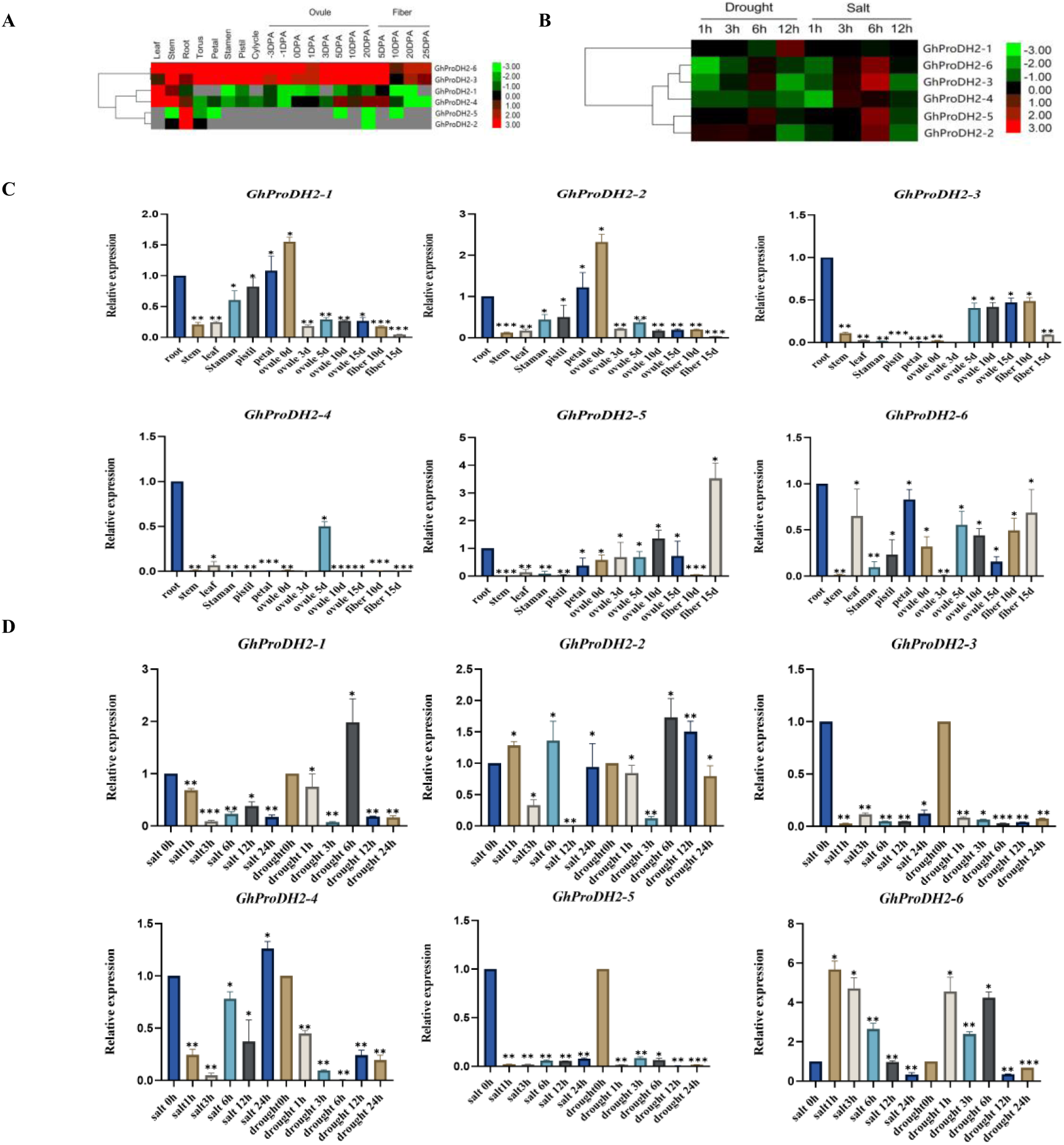
(A) Expression patterns of *GhProDH* genes in various tissues, (B) Expression patterns of *GhProDH* genes under different stresses. (C) Expression patterns of *GhProDH* genes in various tissues, qRT-PCR analysis verifying the expression pattern of *GhProDH* genes in tissues while (D) Expression patterns of *GhProDH* genes under different stresses, qRT-PCR based verification of expression pattern of *GhProDH* gene under stress The error bars represent ± SD of at least three biological replicates (* *P* < 0.05, ** *P*< 0.01, *** *P* < 0.001).

**appendix figure 3.**
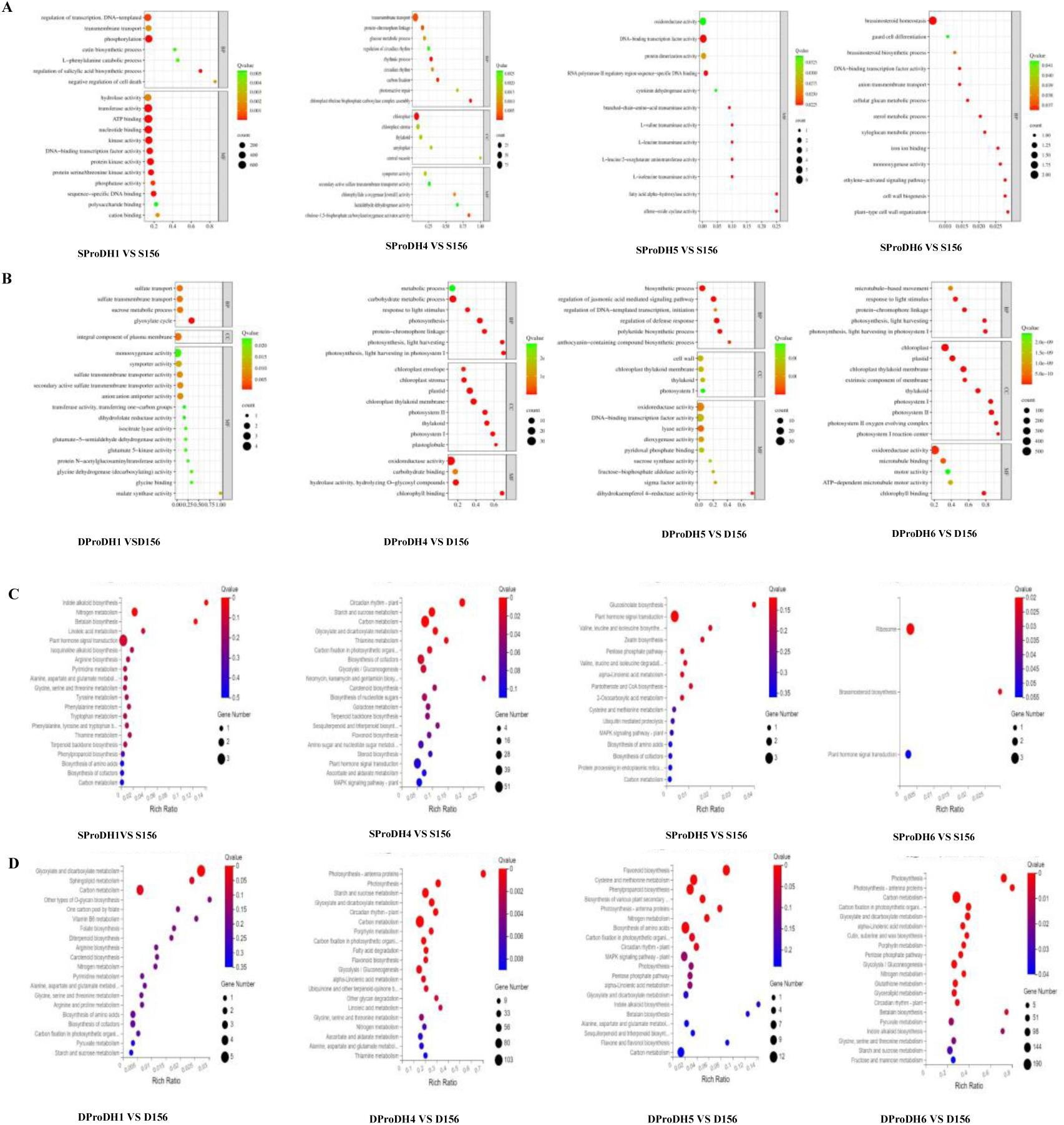
(A) GO enrichment analysis of GhProDH1/156, GhProDH4/156, GhProDH5/156, and GhProDH6/156 under salt stress. (B) GO enrichment analysis of GhProDH1/156, GhProDH4/156, GhProDH5/156, and GhProDH6/156 under drought stress. (C) KEGG enrichment analysis of GhProDH1/156, GhProDH4/156, GhProDH5/156, and GhProDH6/156 under salt stress. (D) KEGG enrichment analysis of GhProDH1/156, GhProDH4/156, GhProDH5/156, and GhProDH6/156 under drought stress. CK156= pYL156; D156= pYL156 drought 8h; S156= pYL156 salt 8h; CKProDH1= pYL156:*GhProH1*; DProDH1= pYL156: *GhProDH1* drought 8h; SProDH1= pYL156: *GhProDH1* salt 8h; CKProDH4= pYL156:*GhProH4*; DProDH4= pYL156: *GhProDH4* drought 8h; SProDH4= pYL156: *GhProDH4* salt 8h; CKProDH5= pYL156:*GhProH5*; DProDH5= pYL156: *GhProDH5* drought 8h; SProDH5= pYL156: *GhProDH5* salt 8h; CKProDH6= pYL156:*GhProH6*; DProDH6= pYL156: *GhProDH6* drought 8h; SProDH6= pYL156: *GhProDH6* salt 8h

**appendix figure 4:**
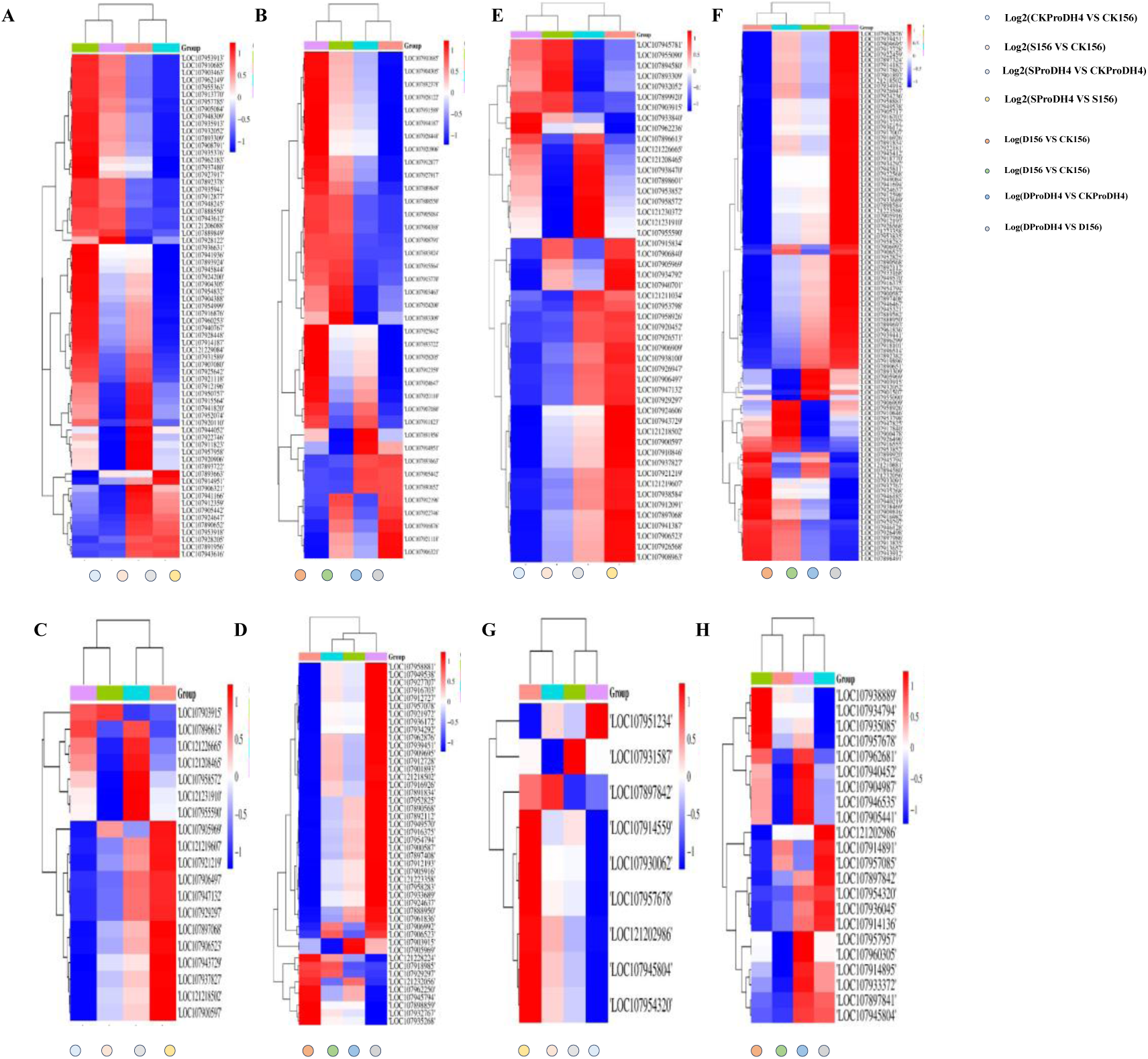
(A) Heat map depicting starch and sucrose pathway clustering for SProDH4/S156, S156/CK156, SProDH4/CKProDH4, and CKProDH4/CK156(B) Heat map illustrating starch and sucrose pathwaglyoxylate dicarboxylic acid pathway clustering for DProDH4/D156, D156/CK156, DProDH4/CKProDH4, and CKProDH4/CK156.(C) Clustering heatmaps illustrating glyoxylate dicarboxylic acid pathway analysis for SProDH4/S156, S156/CK156, SProDH4/CKProDH4, and CKProDH4/CK156. (D) Clustering heatmaps illustrating glyoxylate dicarboxylic acid pathway analysis for DProDH4/D156, D156/CK156, DProDH4/CKProDH4, and CKProDH4/CK156. (E) Heat map illustrating Carbohydrate metabolism for SProDH4/S156, S156/CK156, SProDH4/CKProDH4, and CKProDH4/CK156.(F) Heat map depicting Carbohydrate metabolism for DProDH4/D156, D156/CK156, DProDH4/CKProDH4, and CKProDH4/CK156.(G) Heat map depicting Flavonoid biosynthesis for SProDH4/D156, S156/CK156, SProDH4/CKProDH4, and CKProDH4/CK156. (H) Heat map depicting Flavonoid biosynthesis for DProDH4/D156, D156/CK156, DProDH4/CKProDH4, and CKProDH4/CK156.

**appendix figure 5.**
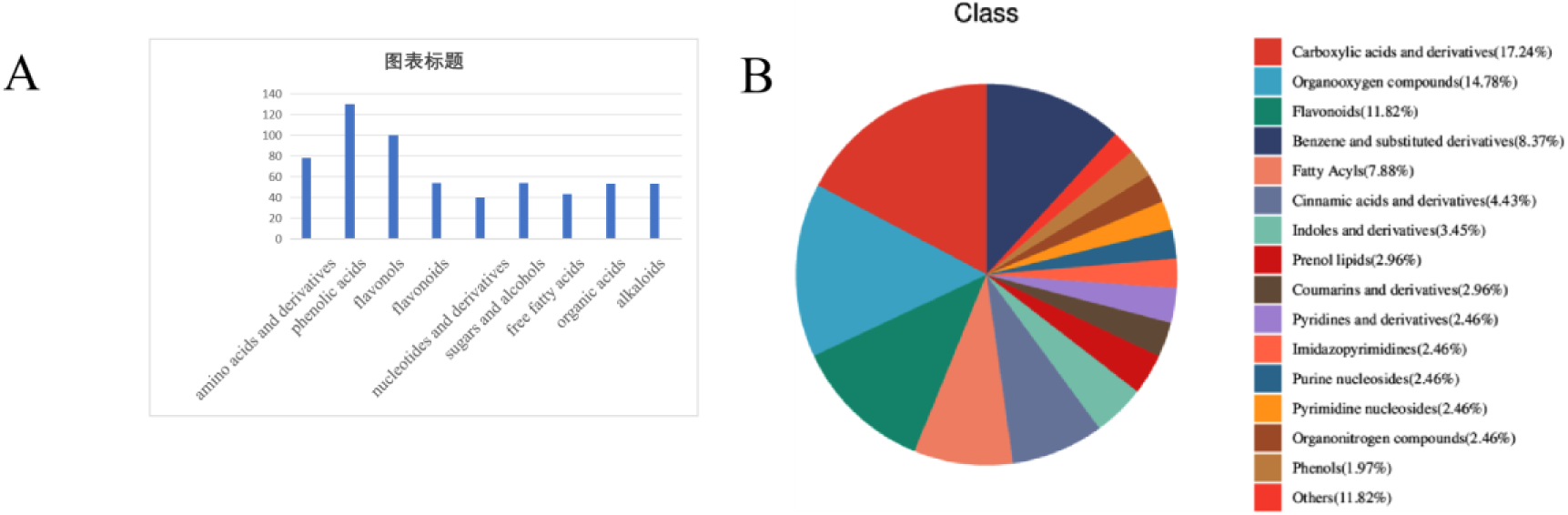
(A) The content of various substances in the metabolome.(B) The proportion of each substance content in the metabolome.

## REFERENCES

Abdelraheem, A., Esmaeili, N., O’Connell, M., and Zhang, J. (2019). Progress and perspective on drought and salt stress tolerance in cotton. Industr. Crops Product. 130, 118–129.

Armengaud, P., L. Thiery, N. Buhot, G. Grenier-De March and A. Savouré (2004). “Transcriptional regulation of proline biosynthesis in Medicago truncatula reveals developmental and environmental specific features.” Physiol Plant 120(3): 442–450.

Baxter, A., R. Mittler and N. Suzuki (2014). “ROS as key players in plant stress signalling.” J Exp Bot 65(5): 1229–1240.

Ben Rejeb, K., C. Abdelly and A. Savouré (2012). “Proline, a multifunctional amino-acid involved in plant adaptation to environmental constraints.” Biol Aujourdhui 206(4): 291–299.

Biancucci, Marco, Roberto Mattioli, Laila Moubayidin, Sabrina Sabatini, Paolo Costantino, and Maurizio Trovato. “Proline Affects the Size of the Root Meristematic Zone in Arabidopsis.” BMC plant biology 15, no. 1 (2015): 263.

Dai, W., M. Wang, X. Gong and J. H. Liu (2018). “The transcription factor FcWRKY40 of Fortunella crassifolia functions positively in salt tolerance through modulation of ion homeostasis and proline biosynthesis by directly regulating SOS2 and P5CS1 homologs.” New Phytol 219(3): 972–989.

Dietrich, K., F. Weltmeier, A. Ehlert, C. Weiste, M. Stahl, K. Harter and W. Dröge-Laser (2011). “Heterodimers of the Arabidopsis transcription factors bZIP1 and bZIP53 reprogram amino acid metabolism during low energy stress.” Plant Cell 23(1): 381–395.

Dos Reis, S. P., Lima, A. M., and De Souza, C. R. B. (2012). Recent molecular advances on downstream plant responses to abiotic stress. Int. J. Mol. Sci. 13, 8628–8647.

Fang, S., Yang, H., Duan, L., Shi, J., & Guo, L. (2023). Potassium fertilizer improves drought stress alleviation potential in sesame by enhancing photosynthesis and hormonal regulation. Plant Physiology and Biochemistry, 200, 107744.

Funck, D., S. Eckard and G. Mueller (2010). “Non-redundant functions of two proline dehydrogenase isoforms in Arabidopsis.” BMC Plant Biology 10(70): (19 April 2010).

Furlan, A. L., E. Bianucci, W. Giordano, S. Castro and D. F. Becker (2020). “Proline metabolic dynamics and implications in drought tolerance of peanut plants.”Plant Physiology and Biochemistry 151: 566–578.

Hanson, J., M. Hanssen, A. Wiese, M. M. Hendriks and S. Smeekens (2008). “The sucrose regulated transcription factor bZIP11 affects amino acid metabolism by regulating the expression of ASPARAGINE SYNTHETASE1 and PROLINE DEHYDROGENASE2.” Plant J 53(6): 935–949.

Hanin, M., C. Ebel, M. Ngom, L. Laplaze and K. Masmoudi (2016). “New Insights on Plant Salt Tolerance Mechanisms and Their Potential Use for Breeding.”Front Plant Sci 7: 1787.

Hoque, M. A., E. Okuma, M. N. Banu, Y. Nakamura, Y. Shimoishi and Y. Murata (2007). “Exogenous proline mitigates the detrimental effects of salt stress more than exogenous betaine by increasing antioxidant enzyme activities.”J Plant Physiol 164(5): 553–561.

Hu, C. A., A. J. Delauney and D. P. Verma (1992). “A bifunctional enzyme (delta 1-pyrroline-5-carboxylate synthetase) catalyzes the first two steps in proline biosynthesis in plants.”Proc Natl Acad Sci U S A 89(19): 9354–9358.

Huang, L.-b., L.-n. Peng and X.-h. Yan (2021). “Multi-omics responses of red algae Pyropia haitanensis to intertidal desiccation during low tides.” Algal research (Amsterdam) 58: 102376.

Huang, Y., Z. Bie, Z. Liu, A. Zhen and W. Wang (2009). “Protective role of proline against salt stress is partially related to the improvement of water status and peroxidase enzyme activity in cucumber [Cucumis sativus].” Soil science and plant nutrition (Tokyo) 55(5): 698–704.

Iqbal, N., S. Umar and N. A. Khan (2015). “Nitrogen availability regulates proline and ethylene production and alleviates salinity stress in mustard (Brassica juncea).”J Plant Physiol 178: 84–91.

Kiyosue, T., Y. Yoshiba, K. YamaguchiShinozaki and K. Shinozaki (1996). “A nuclear gene encoding mitochondrial proline dehydrogenase, an enzyme involved in proline metabolism, is upregulated by proline but downregulated by dehydration in Arabidopsis.” Plant Cell 8(8): 1323–1335.

Krasensky, J. and C. Jonak (2012). “Drought, salt, and temperature stress-induced metabolic rearrangements and regulatory networks.”J Exp Bot 63(4): 1593–1608.

Li, L., Y. Li and G. Ding (2024). “Response mechanism of carbon metabolism of Pinus massoniana to gradient high temperature and drought stress.” BMC Genomics 25(1): 166–166.

Li, Y., D. Jiang, X.-Y. Liu, M. Li, Y.-F. Tang, J. Mi, G.-X. Ren and C.-S. Liu (2023). “Multi-Omics Analysis Provides Crucial Insights into the Drought Adaptation of Glycyrrhiza uralensis Fisch.” J. Agric. Food Chem 71(13): 5391–5402.

Li, W., R. Meng, Y. Liu, S. Chen, J. Jiang, L. Wang, S. Zhao, Z. Wang, W. Fang, F. Chen and Z. Guan (2022). “Heterografted chrysanthemums enhance salt stress tolerance by integrating reactive oxygen species, soluble sugar, and proline.” Hortic Res 9: uhac073.

Liu, C., Deng, Y., Liu, X., & Lin, J. (2023). Research Advances in the Function of Catalase in Plant Growth,Development and Stress Response. Sheng Ming Ke Xue Yan Jiu, (2), 128.

Lv, H., Guo, S., Wu, Z., Nan, X., Zhu, M., & Mao, K. (2024). Postharvest quality and metabolism changes of daylily flower buds treated with hydrogen sulfide during storage. Postharvest Biology and Technology, 212, 112890.

Mattioli, R., Biancucci, M., Lonoce, C., Costantino, P., & Trovato, M. (2012). Proline is required for male gametophyte development in arabidopsis. BMC Plant Biology, 12, 236.

Mattioli, Roberto, Marco Biancucci, Amira El Shall, Luciana Mosca, Paolo Costantino, Dietmar Funck, and Maurizio Trovato. “Proline Synthesis in Developing Microspores Is Required for Pollen Development and Fertility.” BMC plant biology 18, no. 1 (2018): 356.

Meng, Y.-C., Zhang, H.-F., Pan, X.-X., Chen, N., Hu, H.-F., Haq, S. U., Chen, R.-G. (2021). CaDHN3, a Pepper (Capsicum annuum L.) Dehydrin Gene Enhances the Tolerance against Salt and Drought Stresses by Reducing ROS Accumulation. International Journal of Molecular Sciences, 22(6), 3205.

Monteoliva, M. I., Y. S. Rizzi, N. M. Cecchini, M. R. Hajirezaei and M. E. Alvarez (2014). “Context of action of Proline Dehydrogenase (ProDH) in the Hypersensitive Response of Arabidopsis.” Bmc Plant Biology14: 11.

Nanjo, T., M. Kobayashi, Y. Yoshiba, Y. Kakubari, K. Yamaguchi-Shinozaki and K. Shinozaki (1999). “Antisense suppression of proline degradation improves tolerance to freezing and salinity in Arabidopsis thaliana.” FEBS Lett 461(3): 205–210.

O’Brien, M. J., A. Valtat, S. Abiven, M. S. Studer, R. Ong and B. Schmid (2020). “The role of soluble sugars during drought in tropical tree seedlings with contrasting tolerances.”

Per, T. S., Khan, N. A., Reddy, P. S., Masood, A., Hasanuzzaman, M., Khan, M. I. R., & Anjum, N. A. (2017). Approaches in modulating proline metabolism in plants for salt and drought stress tolerance: Phytohormones, mineral nutrients and transgenics. Plant Physiology and Biochemistry, 115, 126–140.

Ren, Y., M. Miao, Y. Meng, J. Cao, T. Fan, J. Yue, F. Xiao, Y. Liu and S. Cao (2018). “DFR1-Mediated Inhibition of Proline Degradation Pathway Regulates Drought and Freezing Tolerance in Arabidopsis.” Cell Rep 23(13): 3960–3974

Rizzi, Y. S., N. M. Cecchini, G. Fabro and M. E. Alvarez (2016). “Differential control and function of Arabidopsis ProDH1 and ProDH2 genes on infection with biotrophic and necrotrophic pathogens.” Molecular Plant Pathology 18(8): 1164–1174.

Roosens, N., T. T. Thu, H. M. Iskandar and M. Jacobs (1998). “Isolation of the ornithine-δ-aminotransferase cDNA and effect of salt stress on its expression in *Arabidopsis thaliana*.” Plant Physiology 117(1): 263–271.

Roosens, N. H., T. T. Thu, H. M. Iskandar and M. Jacobs (1998). “Isolation of the ornithine-delta-aminotransferase cDNA and effect of salt stress on its expression in Arabidopsis thaliana.” Plant Physiol 117(1): 263–271.

Schekaleva, O., Luneva, O., Klimenko, E., Shaliukhina, S., & Breygina, M. (2024). Dynamics of ROS production, SOD, POD and CAT activity during stigma maturation and pollination in Nicotiana tabacum and Lilium longiflorum. Plant Biology (Stuttgart, Germany), Plant biology (Stuttgart, Germany), 2024–09.

Senthil-Kumar, M. and K. S. Mysore (2012). “Ornithine-delta-aminotransferase and proline dehydrogenase genes play a role in non-host disease resistance by regulating pyrroline-5-carboxylate metabolism-induced hypersensitive response.” Plant Cell and Environment 35(7): 1329–1343.

Silveira, J. A., A. Viégas Rde, I. M. da Rocha, A. C. Moreira, A. Moreira Rde and J. T. Oliveira (2003). “Proline accumulation and glutamine synthetase activity are increased by salt-induced proteolysis in cashew leaves.” J Plant Physiol 160(2): 115–123.

Siripornadulsil, S., S. Traina, D. P. S. Verma and R. T. Sayre (2002). “Molecular mechanisms of proline-mediated tolerance to toxic heavy metals in transgenic microalgae.” Plant Cell 14(11): 2837–2847.

Stephens SG, Mosley ME (1974) Early domesticated cottons from archaeological sites in central coastal Peru. Am Antiq 39:109–122

Szabados, L. and A. Savouré (2010). “Proline: a multifunctional amino acid.” Trends in Plant Science 15(2): 89–97.

Tsikas, D. (2017). Assessment of lipid peroxidation by measuring malondialdehyde (MDA) and relatives in biological samples: Analytical and biological challenges. Analytical Biochemistry, 524, 13–30.

Uzilday, B., Turkan, I., Sekmen, A. H., Ozgur, R., & Karakaya, H. C. (2012). Comparison of ROS formation and antioxidant enzymes in Cleome gynandra (C 4) and Cleome spinosa (C 3) under drought stress. Plant Science (Limerick), 182, 59–70

Urmi, T. A., M. M. Islam, K. N. Zumur, M. A. Abedin, M. M. Haque, M. H. Siddiqui, Y. Murata and M. A. Hoque (2023). “Combined Effect of Salicylic Acid and Proline Mitigates Drought Stress in Rice (Oryza sativa L.) through the Modulation of Physiological Attributes and Antioxidant Enzymes.” Antioxidants (Basel) 12(7): 1438.

Van Aken, O., B. T. Zhang, C. Carrie, V. Uggalla, E. Paynter, E. Giraud and J. Whelan (2009). “Defining the Mitochondrial Stress Response in *Arabidopsis thaliana*.” Molecular Plant 2(6): 1310–1324.

Verma, D., S. K. Jalmi, P. K. Bhagat, N. Verma and A. K. Sinha (2020). “A bHLH transcription factor, MYC2, imparts salt intolerance by regulating proline biosynthesis in Arabidopsis.” Febs j 287(12): 2560–2576.

Verslues, P. E. and S. Sharma (2010). ”Proline metabolism and its implications for plant-environment interaction.” The arabidopsis book 8: e0140.

Yang, W., Li, N., Fan, Y., Dong, B., Song, Z., Cao, H., … Fu, Y. (2021). Transcriptome analysis reveals abscisic acid enhancing drought resistance by regulating genes related to flavonoid metabolism in pigeon pea. Environmental and Experimental Botany, 191, 104627.

Wrzaczek, M., M. Brosché and J. Kangasjärvi (2013). “ROS signaling loops - production, perception, regulation.” Curr Opin Plant Biol 16(5): 575–582.

Xu, Y., Zhao, L., Xing, H., Luo, Y., & Wei, Z. (2020). Effects of endophytic bacteria on proline and malondialdehyde of wheat seedlings under salt stress. Shengtai Xuebao = Acta Ecologica Sinica, (11), 3726.

Zeb, A., W. Liu, L. Meng, J. Lian, Q. Wang, Y. Lian, C. Chen and J. Wu (2022). “Effects of polyester microfibers (PMFs) and cadmium on lettuce (Lactuca sativa) and the rhizospheric microbial communities: A study involving physio-biochemical properties and metabolomic profiles.” J Hazard Mater 424(Pt C): 127405–127405.

Zhang, F., J. Huang, H. Guo, C. Yang, Y. Li, S. Shen, C. Zhan, L. Qu, X. Liu, S. Wang, W. Chen and J. Luo (2022). “OsRLCK160 contributes to flavonoid accumulation and UV-B tolerance by regulating OsbZIP48 in rice.” Sci. China Life Sci 65(7): 1380–1394.

Zhang, J., B. Sun, C. Yang, C. Wang, Y. You, G. Zhou, B. Liu, C. Wang, J. Kuai and J. Xie (2022). “A novel composite vegetation index including solar-induced chlorophyll fluorescence for seedling rapeseed net photosynthesis rate retrieval.” Computers and electronics in agriculture 198: 107031.

